# Optoception: perception of optogenetic brain stimulation

**DOI:** 10.1101/2021.04.22.440969

**Authors:** Jorge Luis-Islas, Monica Luna, Benjamin Floran, Ranier Gutierrez

## Abstract

How do animals experience brain manipulations? Optogenetics has allowed us to manipulate selectively and interrogate neural circuits underlying brain function in health and disease. However, it is currently unknown whether mice could perceive arbitrary optogenetic stimulation in addition to their evoked physiological functions. To address this issue, mice were trained to report an optogenetic stimulation as a cue to obtain rewards and avoid punishments. It was found that mice could perceive optogenetic manipulations regardless of the brain area modulated, their rewarding effects, or the stimulation of glutamatergic, GABAergic, and dopaminergic cell types. We named this phenomenon optoception. Using optoception, mice could learn to execute two different sets of instructions based on the laser frequency. Importantly, optoception can occur either activating or silencing a single cell type. Our findings revealed that mice’s brains are capable of “monitoring” their self-activity, albeit indirectly, perhaps via interoception or as a discriminative stimulus, opening a new way to introduce information to the brain and control brain-computer interfaces.

## Introduction

The brain can sense and integrate signals arising from inside the body, monitoring the status of the internal milieu in a process named interoception (Craig, 2002; Khalsa and Lapidus, 2016). Thus, in its most inclusive definition, interoception is the perception of the state of the body (Ceunen et al., 2016). This process which is not restricted to visceral stimuli occurs both consciously and non-consciously and comprises all body organs, including the brain (Craig, 2002; Hölzl et al., 1996). It is well established that optogenetic and electrical stimulation of various areas can produce a cornucopia of behavioral changes (Di Scala et al., 1987; Guo et al., 2015; Mazurek and Schieber, 2017; Romo et al., 1998; Verrier et al., 1975; Wu et al., 2016), and it is normally assumed that the evoked behavioral changes reflect the function of the manipulated circuits (but see (Otchy et al., 2015). However, it is currently unknown whether mice could learn solely from perceptible signals arising from stimulating the brain itself, in addition to their ascribed physiological function. To this aim, several behavioral protocols were designed to test whether mice could use optogenetic brain stimulation as a detectable cue to obtain a reward and avoid punishment.

## Results

### Optogenetic stimulation transiently perturb spiking homeostasis

To characterize whether optogenetic stimulation transiently affects spiking homeostasis (Otchy et al., 2015), this study chose the prefrontal cortex (PFC) because its glutamatergic and GABAergic neurons are major players regulating the balance of neural activity (Courtin et al., 2014). This study hypothesized that optogenetic manipulation of either glutamatergic or GABAergic PFC neurons abruptly perturbed spiking homeostasis, albeit with opposite modulatory signs. To this aim, optrode recordings were performed in freely moving mice while they were optogenetically activated in a frequency scanner test (Prado et al., 2016). Glutamatergic and GABAergic neurons of the PFC were stimulated by using the Thy1-ChR2 mice, which express the channelrhodopsin 2 (ChR2) in pyramidal layer V glutamatergic neurons (Kumar et al., 2013), and the VGAT-ChR2 mice, which express ChR2 in GABAergic^VGAT+^ neurons. Thus, when GABAergic neurons are optogenetically activated, they produce an indirect but extensive inhibition (Babl et al., 2019; Zhao et al., 2011). As predicted, this study found that optogenetic activation of these cells produced an opposite modulatory pattern as the laser frequency increased (**Figures 1A-D;** Prado et al., 2016; Zhao et al., 2011). Briefly, in the Thy1-ChR2 mice, 67% of PFC neurons increased their excitability, and 20% inhibited after stimulation at 20 Hz. In contrast, optogenetic activation of GABAergic neurons in the VGAT-ChR2 mice resulted in 3% of neurons being excited and 50% being inhibited (**Figure 1E**). Notably, both brain manipulations transiently perturbed the spiking homeostasis (Berntson and Khalsa, 2021; Maffei and Fontanini, 2009; Prado et al., 2016), promoting an activation/inhibition imbalance.

**Figure 1.**
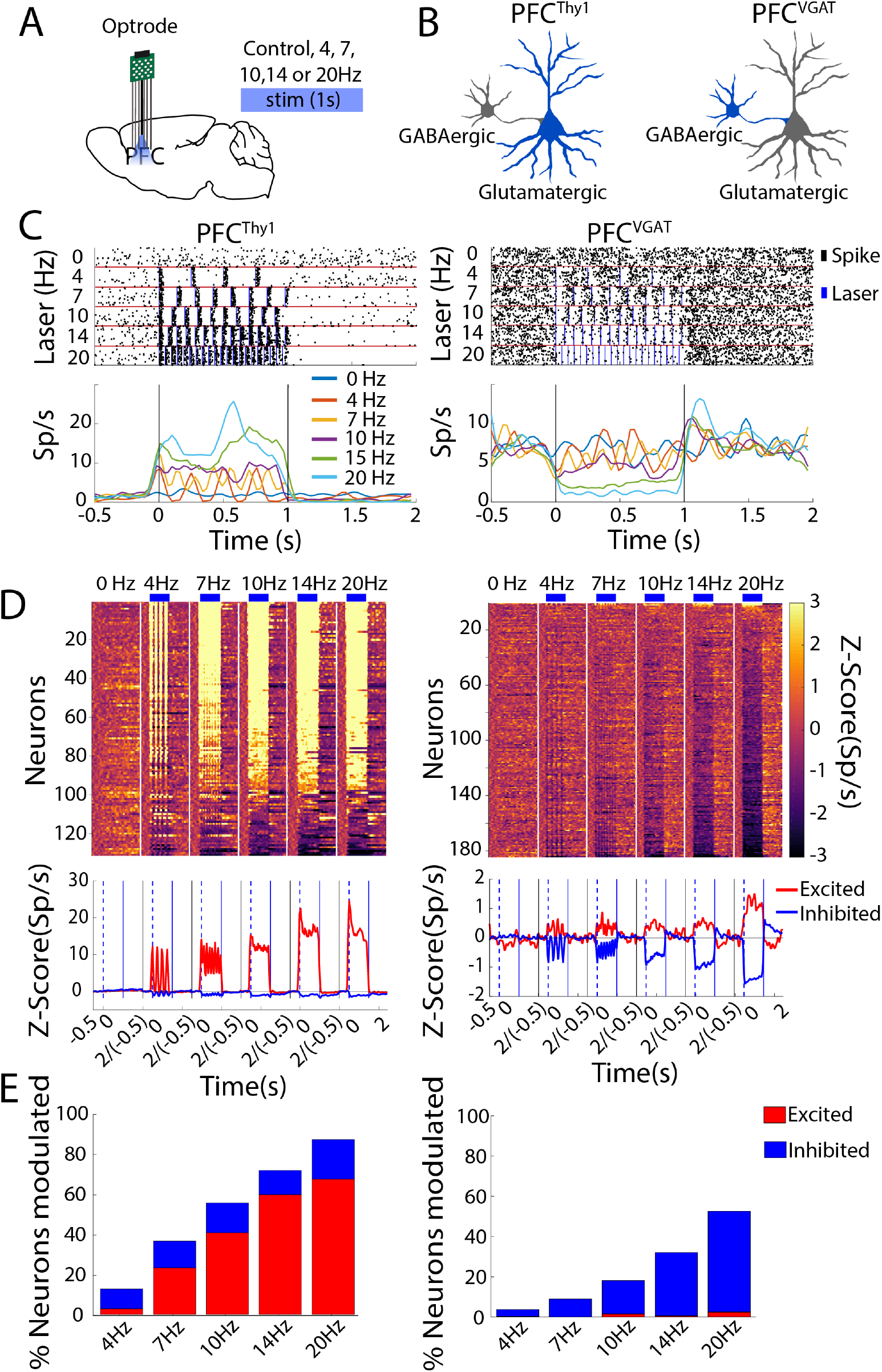
Optogenetic stimulation of PFCThy1 or PFCVGAT transiently impacts spiking excitatory/inhibitory balance, evoking opposite neuronal responses in a laser frequency-dependent manner. (**A**) Schematic representation of PFC recording sites in mice with optrodes record and stimulation at different frequencies. (**B**) Diagram of neurons expressing ChR2 that were optogenetically stimulated (in blue). In PFC^Thy1^ mice, the stimulation drives the activation of glutamatergic neurons, whereas, in PFC^VGAT^ mice, it activates cortical GABAergic neurons inducing an indirect inhibition on glutamatergic neurons. (**C**) Representative examples of two neurons modulated. Upper panel, raster plot of one neuron recorded from PFC^Thy1^ (left) and the other in PFC^VGAT^ (right) aligned to laser onset (Time = 0s). Below are shown the PSTHs, respectively. Vertical lines indicate laser onset. (**D**) Population activity. Upper, a heat map of neuronal population activity from PFC^Thy1^ (left) or PFC^VGAT^ (right), normalized to Z-scores, vertical white lines by laser frequency. Below is the population PSTH activity. Dashed lines indicate laser onset, blue line laser offset, and the black line the trial start. (**E**) Percentage of neurons modulated by different laser frequencies for PFC^Thy1^ (total recorded neurons, n=142, left) or PFC^VGAT^ (total neurons, n= 325, right).

### Mice could perceive an optogenetic perturbation

We then went to test whether, irrespective of cell type or brain region perturbed, optogenetic stimulation induces a stimulus that animals would perceive and learn to use as a cue to obtain rewards and avoid punishment. To test this hypothesis, we optogenetically activated various cell types and brain regions, including the glutamatergic and GABAergic neurons of the PFC. Initially, optical fibers were implanted in the PFC of Thy1-ChR2 mice (PFC^Thy1^), VGAT-ChR2 mice (PFC^VGAT^), or wild-type (WT) mice (PFC^WT^). The WT mice served as a control (**Figure 2A**). The NAc of Thy1-ChR2 mice (NAc^Thy1^) or the Thalamic Reticular Nucleus (TRN) of VGAT-ChR2 mice (TRN^VGAT^) was used to stimulate subcortical brain regions. In the NAc^Thy1^, the glutamatergic afferent fibers innervating the NAc were activated (Prado et al., 2016). In contrast, GABAergic neurons in the TRN of the VGAT-ChR2 mice (TRN^VGAT^) were stimulated (Pinault, 2004). Finally, DA neurons in the Ventral Tegmental Area (VTA) were also optogenetically activated by driving the expression of ChR2 in the TH-Cre mice (VTA^TH^; Gil-Lievana et al., 2020).

**Figure 2.**
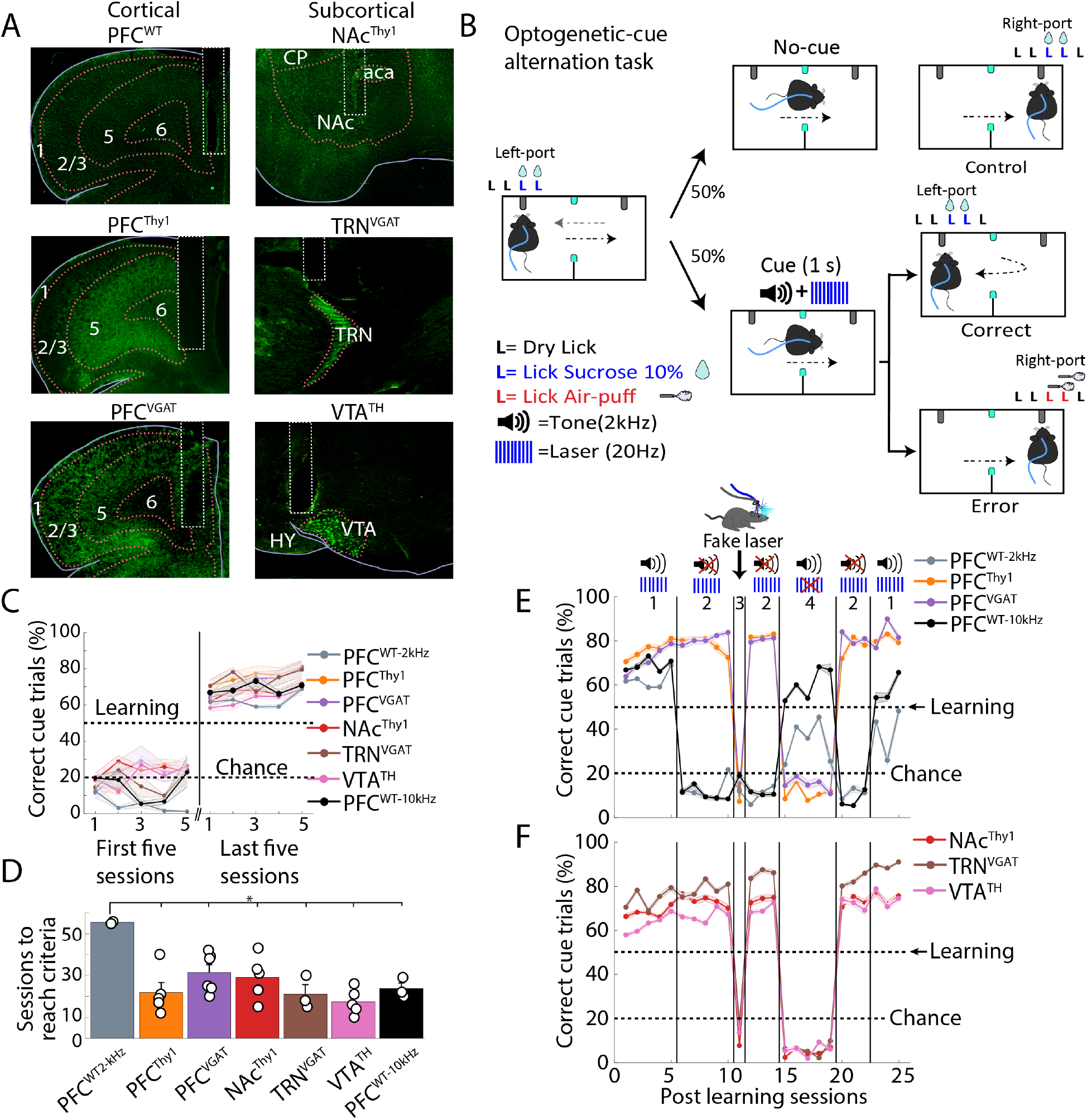
Mice learn to use optogenetic manipulations as a cue regardless of cell type or brain region stimulated. **(A)** Representative images for fiber optics implantation and stimulation sites. *Left pictures* show unilateral optical fibers implanted in prefrontal cortices in wild-type (PFC^WT^), transgenic Thy1-ChR2 (PFC^Thy1^), and VGAT-ChR2 (PFC^VGAT^) mice. *Right pictures* show optical fibers in subcortical regions, including the nucleus accumbens in Thy1-ChR2 (NAc^Thy1^), in the thalamic reticular nucleus in VGAT-ChR2 (TRN^VGAT^), and the ventral tegmental area in TH-Cre (VTA^TH^) mice. (**B**) Schematic of the optogenetic-cue alternation task, where mice had to alternate between two sippers to receive two drops of 10% sucrose from each one. When the mice break the photobeam, located halfway between the two sippers (cyan squares), a cue (tone (2 kHz) + laser (20 Hz), 1s) was randomly delivered. The cue instructed them to return to the previously rewarded port to be rewarded again and avoid punishment. (**C**) Task performance in the initial and last five training sessions. Learning criteria (horizontal dashed line at 50%) was reached when mice avoided punishment in > 50% of cue-trials, in 5 consecutive sessions. Correct trials separated as Hit and Correct Rejections are shown in Figure 2 Supplement 1, where it can be seen that only Hits increased their proportion when mice learned the task. Note that PFC^WT-10kHz^ mice were trained with a more easily perceived auditory tone 10kHz, than a 2 kHz tone that was barely perceptible to mice as shown in Figure 2 Supplement 2. (**D**) Sessions to reach the learning criteria. (**E**) Task performance post-learning in mice with PFC optogenetic stimulation (block 1). In block 2, the tone was removed. In block 3, mice were tested with a “fake laser.” After reacquisition (the second block 2), block 4 began, where the tone was the cue only. Finally, we repeated block 1 (i.e., laser+tone). (**F**) Similar to panel “**E**,” except that stimulation was delivered in subcortical structures (NAc^Thy1^, TRN^VGAT^, and VTA^TH^). \**p*<0.01 ANOVA Dunnet *post hoc* relative to PFC^WT^ control.

To demonstrate that optogenetic stimulation can serve as a discriminative stimulus, initially put to compete with a weak sensory cue (a 2kHz tone) combined with the optogenetic cue. This way, it was possible to show whether animals pay more attention to one cue (and allow WT control mice to learn the optogenetic-cue alternation task). Water-deprived mice were initially habituated to alternate licking between two sippers to obtain sucrose (not shown). After that, they were trained in an optogenetic-cue alternation task, in which 50% of trials (no-cue) continued alternating between sippers to receive sucrose (**Figure 2B**). In contrast, in the other 50% of trials (cue-trials), mice received halfway between sippers a composed cue (tone 2 kHz + laser 20 Hz, 1s), instructing them to stop sipper alternation and return to the previously rewarded sipper to receive sucrose again (Correct cue-trial). If mice ignored the cue, i.e., did not change direction and lick the opposite sipper, two air puffs were delivered as a punishment (**Figure 2B**, Error trial; **Video 1**). We judged that the mice learned the task if, in 5 consecutive sessions, more than 50% of cue-trials were correct (**Figure 2C** and **Figure 2—Supplement 1**). A 2 kHz auditory tone was chosen because it is barely perceptible to mice (Heffner and Heffner, 2007), resulting in a greater saliency towards the optogenetic cue. Of the 10 PFC^WT-2kHz^ mice tested, only 2 solved the task, even though they took significantly more sessions to learn than transgenic mice (**Figure 2D**). The remaining 8 WT mice, in one case even after 130 sessions, never reached the learning criteria (PFC^WT non-L^ trained with a tone 2kHz+laser; **Figure 2—Supplement 2A**). 3 non-learning mice were re-tested by increasing the tone from 2 to 10 kHz (which they readily perceive), and all three mice rapidly reached the learning criterion (**Figures 2C-D** and **Figure 2—Supplement 2B**, PFC^WT-10kHz^). In contrast, all transgenic mice assayed (with tone 2kHz + laser) learned the task (**Figure 2—Supplement 2C-D**), suggesting that mice could perceive optogenetic brain perturbations.

To determine whether transgenic mice used brain manipulations as a discriminative cue, the 2 kHz tone was removed from the combined cue and thus delivered the laser only. As expected, transgenic mice maintained their performance above learning criteria, even when the laser was the only feedback cue (**Figures 2E-F**; see block 2 and **Video 2**). In contrast, PFC^WT^ control mice trained with either 2 or 10 kHz tone dropped their performance at chance level after the tone was removed, demonstrating that WT mice guided their behavior using the tone solely. This result also suggests that WT mice did not use any effect evoked by the blue light laser nor its thermal effects for discrimination (Owen et al., 2019). In sum, unlike controls, all transgenic mice acquire the task, suggesting they used optogenetic stimulation as a discriminative stimulus.

As noted, this study also found that the transgenic mice did not use the blue light *per se* as a cue (Danskin et al., 2015). To further study this in more detail, “fake laser” was used. In this condition, mice could see the blue laser outside the skull without receiving optogenetic stimulation (see **Video 3)**. It was found that the task performance of all transgenic mice dropped at the chance level (**Figures 2E-F**; block 3). Then inquired again whether the tone could also serve as a discriminative stimulus. After reacquisition sessions (only laser; second block 2), the laser was replaced for the tone as a cue (**Figures 2E-F**; block 4). Unlike the WT mice that increased performance with the tone alone, the transgenic mice exhibited a drastic drop in performance (**Video 4**), as observed in the “fake laser” condition. Finally, given that in block 4 and last block 1 (**Figures 2E-F**), the WT mice exhibited a lower overall performance than in the initial block 1, suggest that either WT mice used some information from the combined cue (tone+light) to guide behavior or more likely it reflects a slight extinction induced by the extensive testing with the laser alone (i.e., two consecutive blocks 2). In contrast, our results demonstrate that transgenic mice used optogenetic stimulation to guide behavior and seemed to neglect the tone.

### Mice could learn optoception even when the optogenetic stimulation was the only cue

Importantly and to further demonstrate that transgenic mice do not need an exteroceptive tone to learn, we trained a naïve group of mice but this time only using the laser as a cue. All transgenic mice assayed acquired the task, even when only the optogenetic stimulation served as a discriminative cue (**Figure 3**). These data demonstrate that mice perceive and learn to use optogenetic brain stimulation as a perceptible cue to guide behavior.

**Figure 3.**
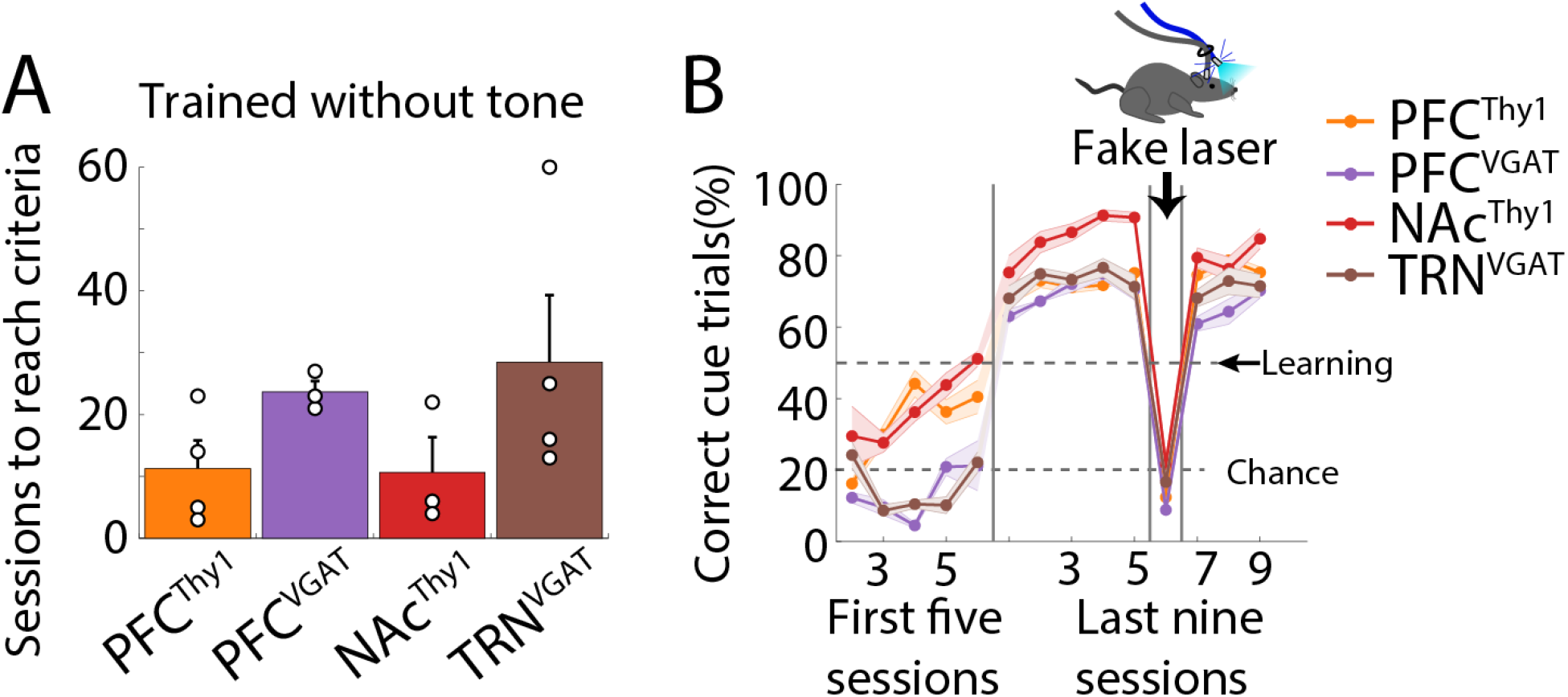
Optoception was induced by optogenetic stimulation alone. (**A**) The number of sessions needed to reach the learning criteria. Mice were trained in optogenetic-cue sipper alternation tasks as shown in **Figure 1B**, but with the laser alone as a cue). Each dot represents an individual subject. (**B**) Percent of correct cue trials. Note the drop in performance during the “fake laser” session, demonstrating that mice used the interoceptive (or any sensory-motor) effects induced by the optogenetic stimulation alone as a conditioned cue.

### Mice could even perceive one single laser pulse

Having demonstrated that mice could use optogenetic stimulation as a cue and since optrode recordings showed that even a single pulse and ≥ 4 Hz laser frequencies could induce a robust neuronal modulation in PFC^Thy1^ and PFC^VGAT^ mice (**Figure 1**). We then characterized the importance of the stimulation parameters to experience optoception, namely frequency and number of pulses. Thus, the same mice in Figure 2 were trained in two variants of the optogenetic-cue alternation task. Consequently, the laser frequency was randomly varied on a trial-by-trial basis (4 to 20 Hz). The correct responses were found to gradually increase as the frequency approached 20 Hz (**Figure 4A**). In the second task variant, the number of pulses was changed (fixed at 20 Hz frequency), and a gradual performance improvement was found when the pulses increased (from 1 to 20). What was somewhat unexpected was that mice could even detect a single pulse (**Figure 4B**). All regions stimulated in these tasks’ variants showed a similar detection profile, except TRN^VGAT^ mice, which were more sensitive and outperformed in both task variants to the other groups. We posit that this arises from its involvement in arousal and attention (Halassa et al., 2014). In sum, it was concluded that mice could also discriminate between different interoceptive (or sensory/motor) stimuli elicited by distinct optogenetic parameters.

**Figure 4.**
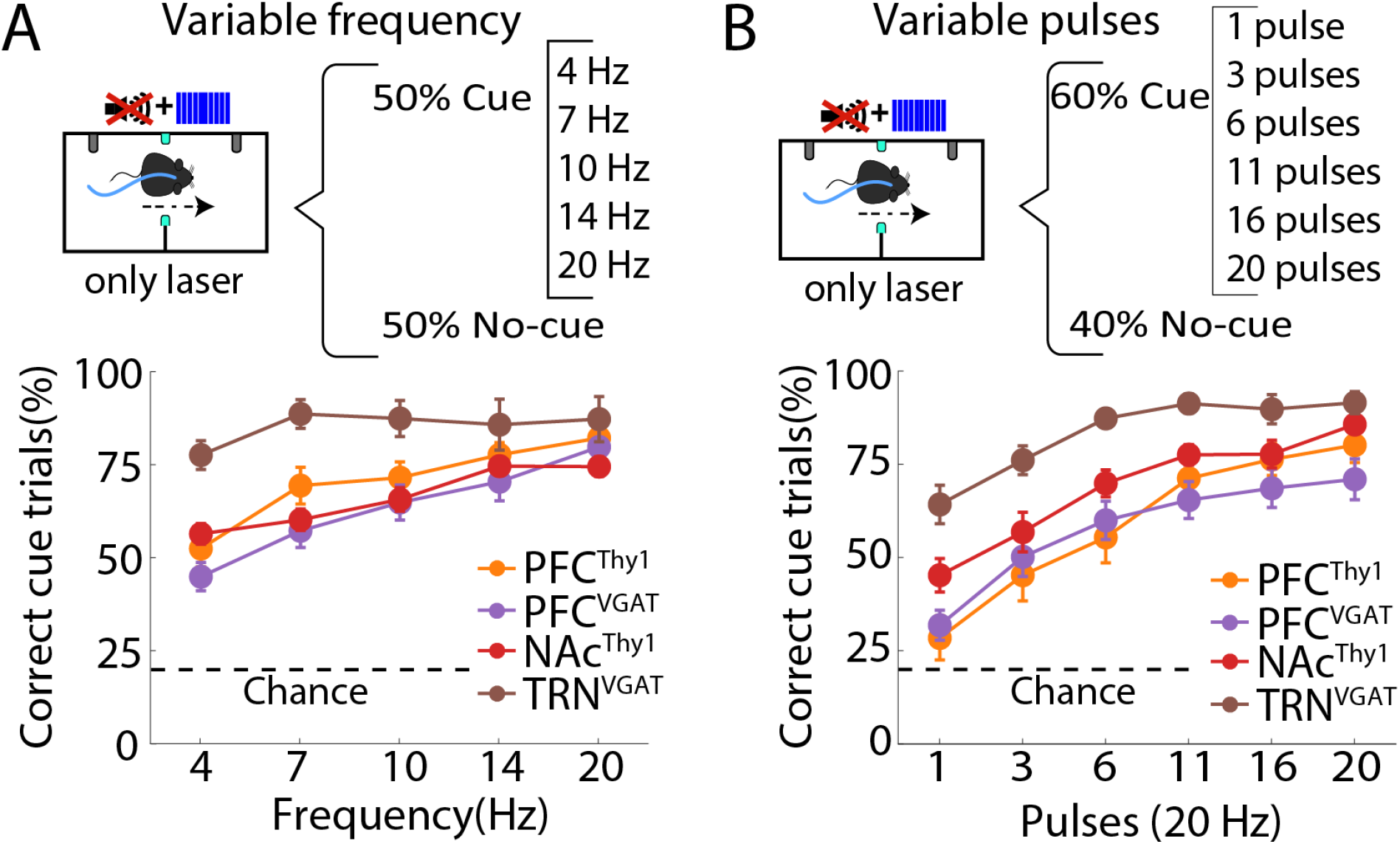
Mice can use optogenetic stimulation only as a cue and generalize to other laser parameters. (**A**) *Upper panel*, schematics of the modified optogenetic-cue alternation task protocol where one out of five frequencies were randomly delivered in 50% of the trials. *The bottom panel* indicates correct cue trials (*correct frequency trials / total frequency trials*). WT mice were not tested in these task variants because they did not perceive the laser alone (see Fig. 1E). (**B**) The *upper panel* depicts the structure of the modified pulse task variant. In this variant, one out of six laser pulses (from 1 to 20) were randomly delivered in 60% of trials. Below is shown the correct cue trials (*correct pulse trials / total pulse trials*). Note that TRN^VGAT^ mice were proficient on both tasks, perhaps because of their role in modulating attention (Wimmer et al., 2015).

### Mice learned two sets of instructions from two different laser frequencies

This study explored whether mice could learn two different sets of instructions based mainly on the laser frequency delivered to the same brain area. In terms of classical perceptual studies, these instruction sets would correspond to two different task paradigms. First, mice were trained in a laser frequency discrimination task (**Figure 5A**), where after visiting the central port, they received either a 10 or 20 Hz stimulation whereupon they had to lick in one of the two lateral ports; one frequency signaled the delivery of sucrose in left port and the other that sucrose is in the right port. If they chose the opposite port, they were punished with two air puffs (**Video 5**). The control PFC^WT^ mice could not learn the task even after 90 training sessions (**Figure 5B**), demonstrating that they did not use the temperature rise elicited by 10 and 20 Hz 1s blue laser stimulation as a discriminative cue (Owen et al., 2019). In contrast, transgenic mice learned this task irrespective of the cell type (glutamatergic and GABAergic) and brain region stimulated, the PFC, NAc, and TRN (**Figure 5B**). All groups learned in a similar number of sessions (one-way ANOVA; F_(3,17)_=2.76, *p* = 0.074). Although the PFC^VGAT^ group exhibited a nonsignificant trend to require additional sessions to reach learning criteria (Holm-Sidak’s multiple comparisons test, *p’s* > 0.05, **Figure 5C)**. These experiments show that transgenic mice can use optogenetic stimulation as a cue. Moreover, in the fake laser session, mice did not use the different light intensity generated from the laser since their performance was at chance level (**Figure 5D**). In a behavioral categorization task, we tested whether mice could categorize laser frequencies. That is, the lower frequencies (10, 12, and 14 Hz) were rewarded if mice went to the left port, and higher frequencies (16, 18, and 20 Hz) were rewarded if they went to the right port (**Figure 5E**; ports were counterbalanced). We found that transgenic mice chose the “high” port more as the laser frequency increased (**Figure 5F**). Thus, mice could also categorize distinct optogenetic laser frequencies.

**Figure 5.**
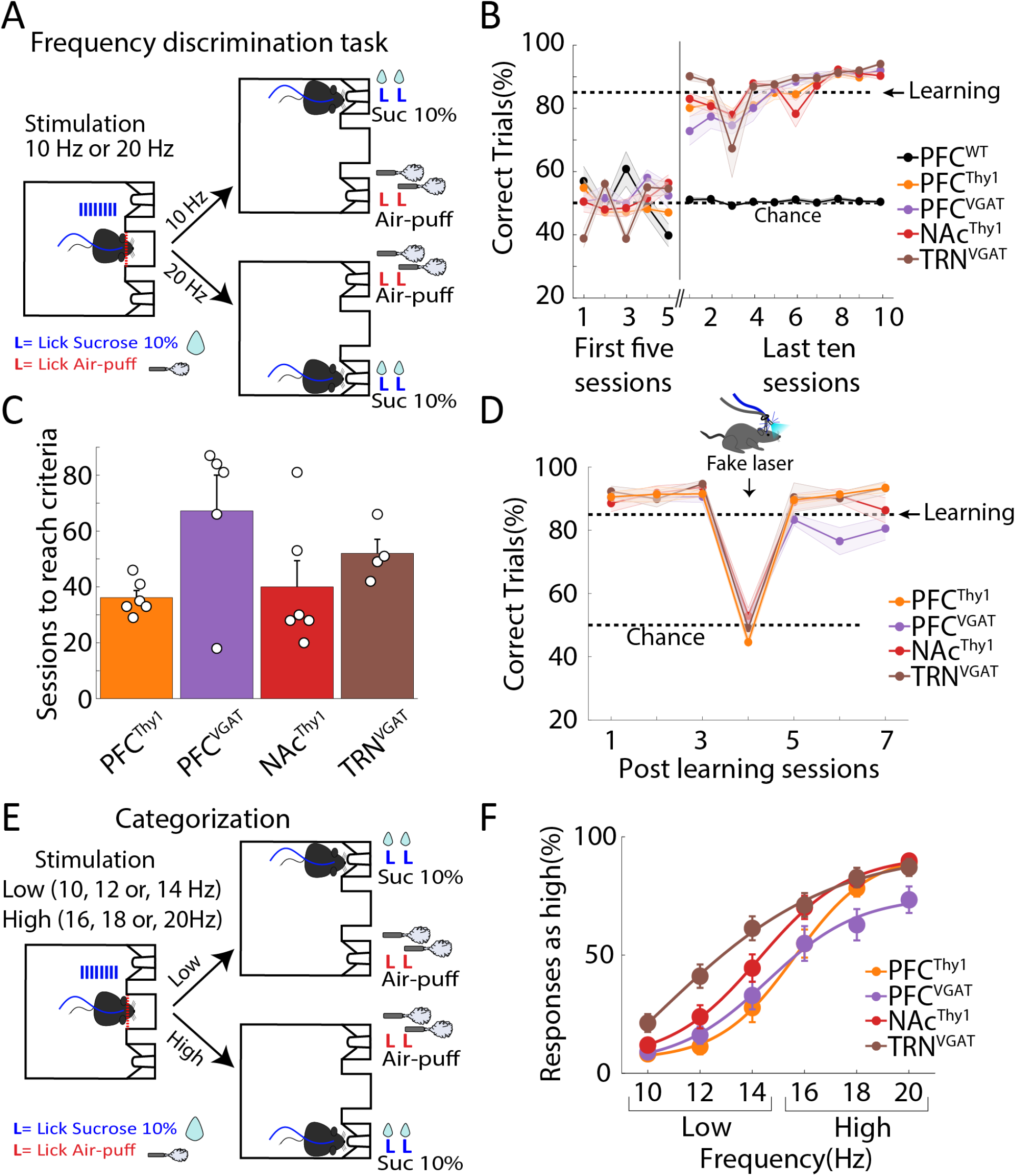
Mice use different laser frequencies to distinguish two actions. **(A)** Scheme of the frequency discrimination task. In this task, upon head entry in the central port (red dashed line), the laser was turned “on” 1 s at 10 or 20 Hz whereupon mice were required to lick in the lateral ports to receive either two drops of sucrose as a reward or two air-puffs as punishment (lateral ports were counterbalanced). (**B**) Correct trials were plotted for the initial five sessions, and the last ten sessions after subjects reached the learning criteria (85% correct trials in 3 consecutive sessions). The control PFC^WT^ mice could not learn even after 90 training sessions. (**C**) The time needed to reach the learning criteria. (**D**) Task performance in subjects that learned the task before and after testing with a “fake laser” in which mice could see the blue light outside the skull but did not receive any optogenetic stimulation. (**E**) Structure of the generalization task, mice had to categorize 10, 12, and 14 Hz frequencies as “low” and 16, 18, and 20 Hz as “high” by licking in the lateral ports. (**F**) Psychometric function for choosing the “high” port. As the laser frequency increases, mice prefer the “high” port more, confirming that they categorized the different laser frequencies. This procedure was counterbalanced across mice.

### Optoception does not require that the optogenetic stimulation be rewarding

It is well-known that rats could guide behavior using the rewarding effects evoked by electrically stimulating the medial forebrain bundle (Wu et al., 2016). Hence, one possibility is that mice learn optoception because the optogenetic modulation rewarded its behavior. For this reason, mice were trained in an operant self-stimulation task to demonstrate that the rewarding effects are important but not essential for optoception. In this task, mice press an active lever to trigger 1s laser stimulation (**Figure 6A and Video 6**). Only PFC^Thy1^ and NAc^Thy1^ mice self-stimulated by pressing the active lever, thereby indicating the stimulation was rewarding (Prado et al., 2016). In contrast, optogenetic stimulation was not rewarding for control PFC^WT^, PFC^VGAT^, and TRN^VGAT^ mice (**Figure 6B**) even though all groups performed equally well in the optogenetic-cue alternation task (**Figure 6C**, see red dots). Again, the rewarding effects of these stimulations were further confirmed in a real-time open field task where mice were optogenetically activated every time they crossed the center of the open field (**Figure 6D and Video 7**). As expected, WT mice rarely visited the center of the open field. However, PFC^Thy1^ somas stimulation and activation of its glutamatergic afferent inputs into the NAc^Thy1^ (Prado et al., 2016) increased the time to cross the center when this zone triggered optogenetic self-stimulation but not during extinction sessions (**Figures 6E-F**). In contrast, stimulation of GABAergic somas in both PFC^VGAT^ and TRN^VGAT^ mice seems to be neutral, i.e., rewarding or aversive effects were not observed (**Figure 6— Supplement 1**). Thus, despite the fact that not all stimulations assayed were rewarding, they all equally served as an optoceptive cue to guide behavior (**Figure 6C**, red dots).

**Figure 6.**
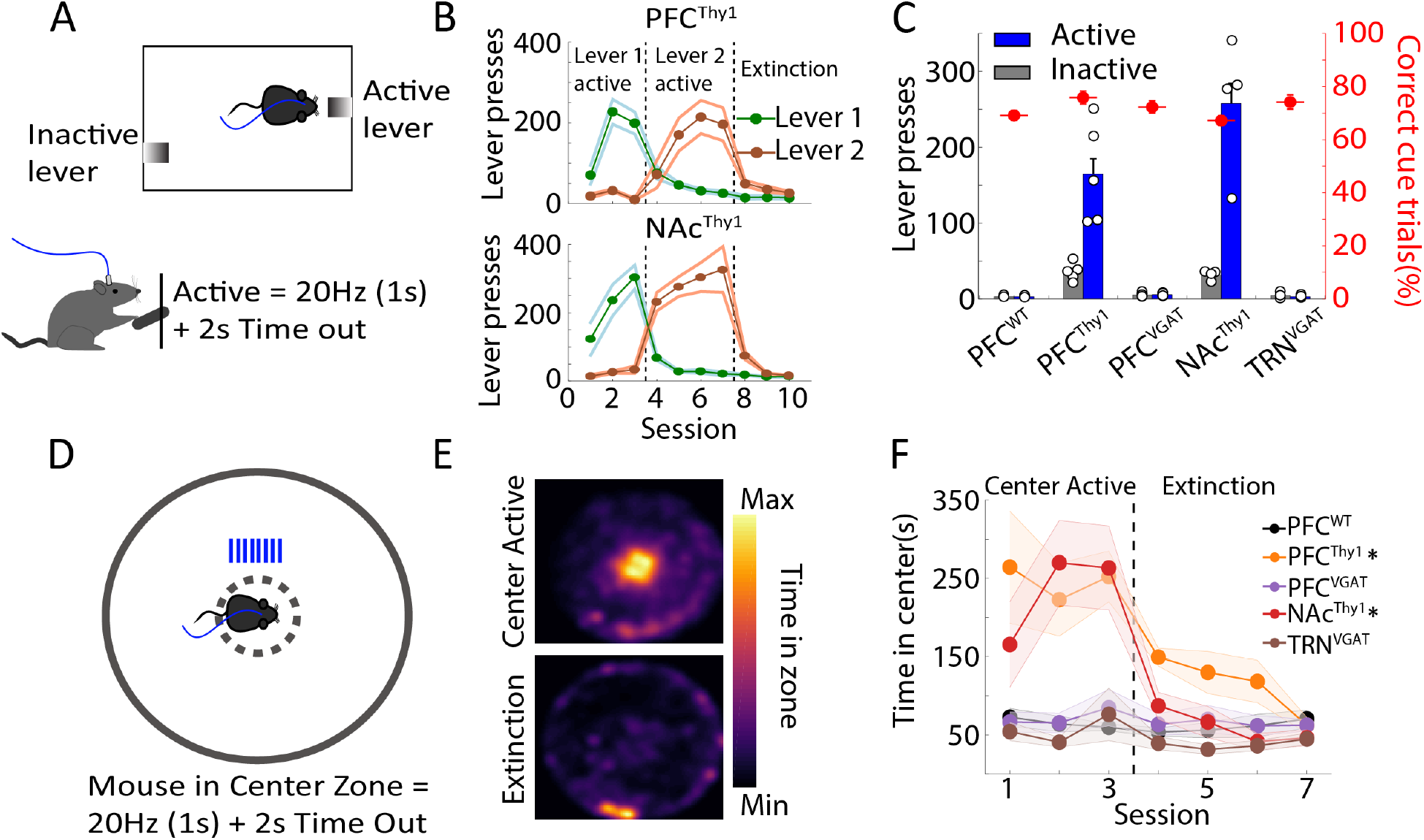
Optoception can guide behavior irrespective of whether brain manipulations elicited rewarding effects or not. (**A**) Scheme of a lever self-stimulation task. Animals can trigger the delivery of laser stimulation by pressing the active lever (20 Hz, 1s + 2s of time out). The inactive lever was recorded but had no programmed consequence. (**B**) The number of lever presses across sessions. This shows that stimulation of PFC^Thy1^ and NAc^Thy1^ was rewarding, as indicated by the number of lever presses. After three sessions, the active lever was switched to inactive and tested for four additional sessions. Levers were counterbalanced across subjects. In the Extinction phase, both levers were Inactive, and thus no laser stimulation was evoked. (**C**) Mean lever presses (excluding Extinction sessions). Small white dots indicate the number of mice tested. Overlapped also shows their average performance (right axis) achieved in the optogenetic-cue alternation task (see solid red circles, Fig. 2C). (**D**) Open field center self-stimulation task. In this task, mice had to cross the center zone to receive laser stimulation (20 Hz, 1s + 2s of time out), note that no other reward or stimuli were delivered. (**E**) Representative heat map of a PFC^Thy1^ mouse that crosses the center (Active) to self-stimulate. The bottom panel shows an extinction session of the same mouse. (**F**) The time spent in the center zone across sessions for all groups. \**p*<0.05, two-way ANOVA, Dunnet *post hoc*, significantly differ from PFC^WT^ during active sessions. Figure 6—Supplement 1. shows that stimulation in PFC^VGAT^ or TRN^VGAT^ mice is not aversive.

### Activating or silencing a single cell type both serve as optoceptive cue

In another test to determine if mice could perceive optogenetic stimuli, we explored if they could learn optoception from activating or silencing a single cell type. This hypothesis was tested using the Vgat-ires-cre mice to drive the selective expression of ChR2 or Archaerhodopsin (ArchT) in GABAergic neurons in the Lateral Hypothalamus (**Figure 7A**; LH^ChR2^ and LH^ArchT^, respectively). Thus, we could activate LH GABAergic neurons with ChR2 or silence them with the outward proton pump, ArchT (Chow et al., 2010). It was found that these mouse types could learn to use both the optogenetic activation and silencing as a cue to solve the optogenetic-cue alternation task (**Figure 7B**). However, stimulating LH GABAergic neurons induced a faster (**Figure 7B**; unpaired t-test, t_(11)_= 3.774, *p*<0.01) and better performance than silencing them (**Figure 7C**; two-way ANOVA, interaction mice x blocks, F_(6,305)_=35.4, *p*<0.0001). Moreover, these mice also maintained their performance above chance level once the tone 2kHz was removed from the combined cue (i.e., they received the laser alone; **Figure 7C**; block 2). In contrast, their performance dropped to chance level when they were tested with a “fake laser” (**Figure 7C**; block 3) or when only the tone was delivered as a cue (**Figure 7C**; block 4). Taken together, these results proved that these mice neglected the 2kHz tone and used the interoceptive (or any sensory-motor) stimuli induced by optogenetic manipulations to guide behavior. LH GABAergic neurons were chosen because it is well established that they could cause opposing behavioral effects (Garcia et al., 2021; Nieh et al., 2016). For example, in a real-time conditioned place preference task (rtCPP), we corroborated that soma stimulation of GABAergic neurons (LH^ChR2^) is rewarding (**Figures 7D-E**) in the sense that they preferred the side paired with the laser stimulation, whereas silencing GABAergic neurons (LH^ArchT^) was aversive since the mice avoided the side paired with the laser (**Figures 7D-E**). Opposing effects on feeding behavior were also observed in LH neurons (**Figures 7F-G**). That is, in sated mice, stimulation of LH GABAergic neurons promoted consumption (**Figures 7F**; LH^ChR2^), whereas in water-deprived mice, silencing them reduced sucrose intake (**Figures 7G** in LH^ArchT^, and **Figure 7—Supplement 1**). In sum, although activating (or silencing) LH GABAergic neurons had opposing effects on reward and feeding, both manipulations were perceived and used as feedback cues to guide behavior.

**Figure 7.**
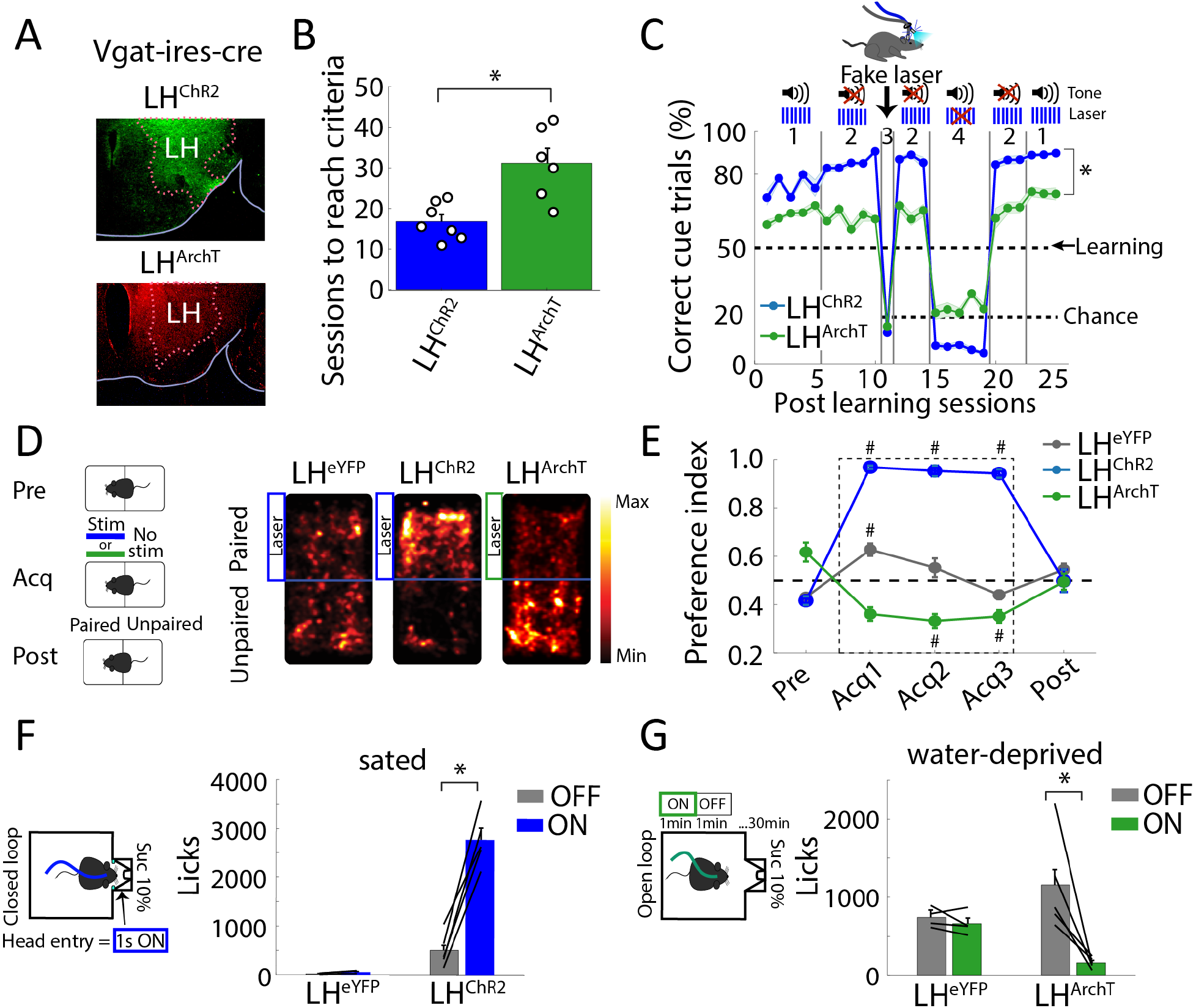
Mice could use both activation or silencing of a single cell type as a perceptible cue, even though they evoked opposing behavioral effects upon reward and feeding. (**A**) Histology of mice transfected with ChR2 or ArchT in Vgat-ires-cre mice (GABAergic neurons) of the Lateral Hypothalamus (LH^ChR2^ or LH^ArchT^, respectively). (**B**) Sessions to reach learning criteria. Each dot represents a mouse. \**p*<0.001 unpaired t-tests. (**C**) Correct trials in the presence of tone (2kHz) and/or laser. Same conventions as in Figure 2e. \**p*<0.001 ANOVA two-way (transgenic mice x block). (**D**) Real-time conditioned place preference (rtCPP). Left, rtCPP task consisted of three phases. Pre-test (Pre, 1-session), Acquisition (Acq, 3-sessions), and Post-test (Post, 1-session). Right, representative heat maps on the acquisition phase. Transfected Vgat-ires-cre mice with the enhanced Yellow Fluorescent Protein (LH^eYFP^) were used as control (**E**) Fraction of time spent on the paired side. Stimulation in LH^ChR2^ mice was rewarding (value>0.5) while silencing in LH^ArchT^ was aversive (<0.5). #*p*<0.0001, ANOVA Dunnett *post hoc*, relative to pre-test. (**F**) Left, schematic of the closed-loop task. Sated LH^ChR2^ or LH^eYFP^ mice were placed in a behavioral box with a sucrose sipper. Head entry into the port triggered optogenetic stimulation (1 s “on,” 20 Hz + 2 s timeout, 473nm). Right, total licks during the task. (**G**) Left panel, open-loop task. In water-deprived LH^ArchT^ or control LH^eYFP^ mice, a continuous green laser was turned “on” in blocks of 1 min (532 nm) and 1 min with no-laser (“off”). Right, total licks during the task. \**p*<0.001 paired t-test. Figure 7—Supplement 1 depicts a raster plot of sucrose licking during stimulation of LH^ChR2^, LH^ArchT^, and LH^eYFP^ mice.

## Discussion

The ability to manipulate the activity of genetically defined cell types via optogenetics has been a game-changing technology in neuroscience. However, very little is known about their sensory-evoked effects. Our findings collectively reveal that mice were capable of perceiving arbitrary optogenetic stimulation. It was found that mice detected and actively reported activation and silencing of various cell types and brain regions. Mice could even sense a single laser pulse, discriminate, and categorize between distinct laser frequencies. Moreover, optoception occurred even when optogenetic activation or silencing of a single cell-type elicited rewarding or aversive effects or whether it promoted feeding or stopped it, respectively. It was proposed that mice perceived the interoceptive state evoked by the optogenetically activated brain circuit and then learned to use it as a conditioned cue to guide behavior. This aligns well with the findings of Doty 1965, who trained monkeys to report electrical stimulation by pressing a lever to obtain a reward and avoid an electric shock on the leg or tail (Doty, 1965), as well as those of Mazurek & Shieber 2017 who asked them to discriminate the parameter of intracranial stimulation delivered in premotor cortex (Mazurek and Schieber, 2017). In both cases, monkeys could detect when and whereinto the premotor cortex of the stimulation was delivered. Our results further extend these observations to optogenetic manipulations most often employed. In sum, this study proposed that if optogenetic activation is strong enough to perturb spiking homeostasis, then they will induce an interoceptive signal (or any sensory-motor discriminative stimulus) that would make mice aware of the brain region that was either activated or silenced optogenetically (Berntson and Khalsa, 2021). Given that the primary goal of optogenetics is to determine the physiological function (i.e., necessity and sufficiency) that a given cell type supports, this study’s results revealed an unexpected but important side effect evoked by the most commonly performed optogenetic perturbations. Thus, more attention should be paid to the perceptible effects induced by optogenetics.

How humans and mice experience brain manipulations -electrical or optogenetic- is an intriguing question (Mazurek and Schieber, 2019). Are they experienced as a natural or artificial stimulus? One can argue that co-activation of an arbitrarily large number of neurons rarely occurs under physiological conditions (with optogenetics, this effect is perhaps exacerbated since only one specific cell type and associated brain circuits are co-activated). Thus, it would be most likely experienced as artificial (Mazurek and Schieber, 2019). However, more local intracranial microstimulation of the somatosensory cortex (S1) seems to be experienced more naturally since it could even be substituted for a natural sensory stimulus (Romo et al., 1998; Tabot et al., 2013). Of course, those studies could not rule out some degree of embodiment (Mazurek and Schieber, 2019). The seminal work of Wilder Penfield revealed that electrical stimulation could induce a “brainbow” of effects comprising noticeable movements, urge to move, somatosensory, visual, or auditory percepts, skin tingling or numbness, as well as rewarding or aversive effects, and even complex emotions (Penfield and Rasmussen, 1950). However, other instances produced no identifiable effect (Penfield and Rasmussen, 1950). Furthermore, various cell type optogenetic manipulation of the S1 cortex bias (i.e., potentiate or suppress) somatosensory perception, whereas others did not affect perception at all (Sun et al., 2021). Based on these results, in our experiments, it would be unlikely that mice felt precisely the same interoceptive (or any other sensory) sensation in each optogenetic manipulation assayed herein. What is clear is that all transgenic mice acquired the task. However, some subjects (or stimulation sites) differed in performance and required more sessions to achieve learning, suggesting that not all optogenetic manipulations were equally perceived. Likewise, our data do not rule out the possibility that some brain regions, or less intense stimulation parameters, could not be perceived at all. Nevertheless, our results demonstrate that some optogenetic manipulations are perceived. Thus, they may also be a helpful animal model to investigate specific interoceptive states evoked by various cell types (Berntson and Khalsa, 2021), akin to the interoceptive conditioning phenomenon (Razran, 1961) elicited by drug-induced body states (Bevins and Besheer, 2014; Ceunen et al., 2016; Solinas et al., 2006). Nevertheless, our results suggest that the rewarding, neutral, or aversive effects induced by optogenetic manipulations are unnecessary to experience optoception since mice learn to use all three interoceptive states as a discriminative stimulus. We posit that mice would be aware of most, if not all, optogenetic brain stimulation, probably using interoception (or any other sensory-motor stimuli evoked by the stimulation).

This study’s results demonstrate that the brain is capable of “monitoring” its self-activity (via its evoked interoceptive state or discriminative stimuli), as previously suggested but not demonstrated by classic experiments of volitional control of neural signals proposed by Eberhard E. Fetz (Fetz, 1969), since these experiments necessarily require an exteroceptive sensory stimulus (auditory or visual) as a feedback cue to learn (Koralek et al., 2012).

The use of optoception also implies that it can be implemented as an independent sensory channel to control brain-computer interfaces. This idea has been recently demonstrated by Prsa et al. 2017 who showed that mice could use artificial optogenetic stimulation of the S1 cortex as sensory feedback to accelerate the control of their M1 neuronal activity (Prsa et al., 2017), thereby presenting an opportunity for using optoception as a parallel information channel to perform brain-computer interfaces (Koralek et al., 2012; Yadav et al., 2021). Our results extend these observations to show that the cortex or subcortical regions and stimulating or silencing a single cell type could be used as an additional sensory channel to introduce information to the brain.

Finally, given that the primary goal of optogenetics is to unveil the function of a given cell type, our results highlight the importance of considering their perceptible effects in interpreting optogenetic experiments, given that mice could also learn from brain stimulation *per se*.

## Material and Methods

### EXPERIMENTAL MODEL AND SUBJECT DETAILS

We used adult male and female mice weighing 20-30 g at the beginning of the experiment. Different strains of mice were used in this study C57Bl6J (WT mice), B6.Cg-Tg(Thy1-COP4/eYFP)18Gfng/J (hereafter referred to as *Thy1-ChR2 mice*), B6.Cg-Tg(Slc32a1-COP4*H134R/eYFP)8Gfng/J (*VGAT-ChR2 mice)*, mice were purchased from the Jackson Laboratory (RRID : IMSR_JAX:007612 and RRID : IMSR_JAX:014548, respectively). The Thy1-ChR2 mice expressed the light-activated ion channel, Channelrhodopsin-2 (ChR2), fused to an enhanced Yellow Fluorescent Protein (eYFP) under the control of the mouse thymus cell antigen 1 (Thy1) promoter (Kumar et al., 2013). The VGAT-ChR2 mice expressed the ChR2 under the control of vesicular GABA transporter (VGAT; Zhao et al., 2011). Furthermore, we employed the Vgat-ires-cre mice because they express cre-recombinase enzyme under the control of endogenous Vgat promoter (RRID:IMSR_JAX:016962; Vong et al., 2011). Finally, we used TH-Cre mouse, which expresses cre-recombinase under the control of endogenous tyrosine hydroxylase promoter. These transgenic mice were kindly donated by Dr. Bermúdez-Rattoni from Instituto de Fisiología Celular, Universidad Nacional Autónoma de México.

Mice were individually housed in standard laboratory cages in a temperature-controlled (22 ± 1°C) room with a 12:12 h light-dark cycle (lights were on 07:00 and off at 19:00). All procedures were approved by the CINVESTAV Institutional Animal Care and Use Committee. Unless otherwise stated, mice were given *ad libitum* access to water for 60 min after testing. Chow food (PicoLab Rodent Diet 20, St. Louis, MO, USA) was always available in their homecage. All experiments were performed in the light period from 09:00 to 16:00h

### METHOD DETAILS

#### Viral constructs

The Cre-inducible adeno-associated virus (AAV) were purchased from *addgene*. For ChR2-eYFP (AAV5-EfIa-DIO-hChR2(E123T/T159C)-EYFP, #35509, at titer of 1×10^13^ vector genome/ml (vg/ml), ArchT-tdTomato (AAV5/FLEX-ArchT-tdTomato), #28305, at titer of 7.0×10^12^ vg/ml and eYFP-vector (AA5V-EfIa-DIO EYFP), #27056, at titer of 1.0×10^13^ vg/ml. Viruses were subdivided into aliquots and stored at −80 °C before their use.

#### Stereotaxic surgery

Mice were anesthetized with an intraperitoneal injection of Ketamine/Xylazine (100/8 mg/Kg). Then mice were put in a stereotaxic apparatus where a midline sagittal scalp incision was made to expose the skull to insert two holding screws.

##### Viral infection

A microinjection needle (30-G) was connected to a 10 μl Hamilton syringe and filled with adeno-associated virus (AAV). Vgat-ires-cre or TH-cre mice were microinjected with AAV (0.5 μl) at a rate of 0.2 µl/min. The injector was left in position for 5 additional minutes to allow complete diffusion. In the case of Vgat-ires-cre, the microinjection was performed unilaterally in the Lateral Hypothalamus (LH, from bregma(mm): AP -1.3, ML ±1.0 and from dura (mm): DV -5.5) for the expression of ChR2 (LH^ChR2^), ArchT (LH^ArchT^) or eYFP (LH^eYFP^), and then a zirconia ferrule of 1.25 mm diameter with multimode optical fibers (200 μm, Thorlabs) was implanted in LH (from bregma(mm): AP -1.3, ML ±1.0 and from dura(mm): DV -5.3). TH-Cre mice were microinjected bilaterally with ChR2 in VTA (TH-VTA mice; from bregma(mm): AP -3.0, ML ±0.6, and DV -4.8), and ferrules were implanted in VTA (from bregma(mm): AP -3.0, ML ±1.2, and DV -4.3, 10° angle), mice were allowed one month for recovery and obtain a stable expression of ChR2 or ArchT.

##### Fiber optics and optrode implantation

A Unilateral zirconia ferrule was implanted in WT, Thy1-ChR2, and VGAT-ChR2 mice. Thy1-ChR2 mice were implanted in prefrontal cortex (*PFC*^*Thy1*^, from bregma(mm): AP +1.94, ML ±0.3, and DV -2.8), or in Nucleus Accumbens (NAc^Thy1^, from bregma(mm): AP +1.2, ML ±1.0, and DV -5.2). For VGAT-ChR2 mice, the optical fiber was implanted in PFC under the same coordinated as the *PFC*^*Thy1*^ (PFC^VGAT^) or in Thalamic Reticular Nucleus (TRN^VGAT^, from bregma(mm): AP -1.55, ML ±2.5, and DV -3.25). WT mice were implanted in the PFC under the same coordinates as the *PFC*^*Thy1*^ (PFC^WT^). The optrode comprised a homemade array of 16 tungsten wires (35 μm) formvar insulated (California Fine Wire Company), surrounded in a circular configuration a single optical fiber was implanted in PFC in Thy1-ChR2 (n=3) or VGAT-ChR2 (n=3) mice (from dura (in mm): AP +1.94, ML ±0.3 and DV -2.8). Mice were allowed one week for recovery from surgery.

#### Optogenetic parameters

Mice expressing ChR2 were stimulated with a diode-pumped solid-state system blue at 473 nm (OEM laser, UT, USA) or green at 532 nm for ArchT opsin (Laserglow Technologies, Toronto, Canada). The light output intensity at the fiber optic patch cord for ChR2 was 3 mW for *PFC*^*Thy1*^, whereas 15 mW for the remaining transgenic mice, while for ArchT stimulation, it was 20 mW. In control *PFC*^*WT*^ mice, they were photo-stimulated at 3mW in the optogenetic-cue alternation task and 15 mW for the frequency discrimination task. It was measured with a fiber optic power meter with an internal sensor (PM20A, Thorlabs, NJ, USA). The pulse lengths were 30 ms for ChR2 and continuous pulse for ArchT and were controlled by Med Associates Inc, software, and TTL signal generator (Med Associates Inc., VT, USA).

#### Extracellular optrode recordings

Thy1-PFC and VGAT-PFC mice with optrode implant were recorded for 20 min in a scanner laser frequency task (Prado et al., 2016), where they received a random stimulation frequency during 1s “on,” followed by 2s “off,” and the subsequent frequency was randomly chosen. The stimulation frequencies were Control (0 Hz), 4, 7, 10, 14, and 20 Hz. Mice were recorded in a maximum of 7 consecutive sessions (**Figure 1**).

Neural activity was recorded using a Multichannel Acquisition Processor system (Plexon, Dallas, TX) interfaced with a Med Associates conditioning side to record behavioral events simultaneously. Extracellular voltage signals were first amplified x1 by an analog headstage (Plexon HST/16o25-GEN2-18P-2GP-G1), then amplified (x1000) sampled at 40 kHz. Raw signals were band-pass filtered from 154 Hz to 8.8 kHz and digitalized at 12 bits resolution. Only single neurons with action potentials with a signal-to-noise ratio of 3:1 were analyzed. The action potentials were isolated online using voltage-time threshold windows and three principal components contour templates algorithm. A cluster of waveforms was assigned to a single unit if two criteria were met: Inter-Spike Intervals were larger than the refractory period set to 1 ms, and if a visible ellipsoid cloud composed of the 3-D projections of the first three principal component analysis of spike waveform shapes was formed. Spikes were sorted using Offline Sorter software (Plexon, Dallas, TX). Only time stamps from offline-sorted waveforms were analyzed. All electrophysiological data were analyzed using MATLAB (The MathWorks Inc., Natick, MA; (Fonseca et al., 2018).

#### Behavioral methods

##### Habituation phase: Training to alternate between sippers

Mice were initially trained in a behavioral box (31 × 41.5cm) that was equipped with two sippers at each end of the box (**Figure 2B**). Each sipper was calibrated to deliver two drops of sucrose 10% (3-4 μl each drop) or two air-puffs (10 psi) via a computer-controlled solenoid valve (Parker, Ohio, USA). The two sippers were semi-divided by a central acrylic wall covering 2/3 parts. This forced mice to cross between sippers by the remaining 1/3 part of the wall that remains open, facilitating the detection of mice’s crosses using a photobeam in the central wall that was recorded when mice were halfway between the two sippers.

Mice were water-deprived for 23h and then placed in the behavioral box. In the first phase, they had to learn to alternate between the two sippers. In each trial, mice had to lick an empty sipper twice to receive two sucrose drops in the third and fourth lick (trial). They need to move to the opposite sipper to start a new trial and obtain more sucrose. Each session lasted 30 min daily. To consider that a mouse learned, they had to complete 60 trials in one session.

##### Optogenetic-cue alternation task

Once mice learned to alternate between sippers, they were placed in a similar task, but this time in 50% of trials, a cue was delivered randomly. This consisted of a tone 2 kHz + laser at 20Hz (or continuous pulse for ArchT) delivered when mice were heading halfway toward the opposite sipper; at this moment, the mice had to return to the previous trial’s sipper to initiate a new rewarded trial. If they licked on the opposite sipper, the mice were punished with 2 air-puffs delivered in the third and fourth lick (**Figure 2B**). The mice were trained on this task until they reached 5 consecutive sessions with >50% of punishments avoided. The task consisted of four different blocks, which differ in the type of cue delivered. In block 1, the combined tone and optogenetic stimulation served as a cue. In block 2, the optogenetic stimulation alone was used as a cue, whereas in block 3, a “fake laser” was delivered. The fake laser was used only as a visual cue. A fiber optic was attached outside the skull (not connected to the implant), and thus it did not provide an optogenetic stimulation. Finally, block 4 used only the auditory cue. All sessions were 30 min long.

##### Frequencies and pulse task variants

The same mice were also tested in 5 consecutive sessions in the optogenetic-cue alternation task. The laser frequency changed trial by trial in random order at 4, 7, 10, 14, and 20 Hz in 50% of total trials (**Figure 4A**). In the subsequent 5 sessions, the laser frequency was fixed constant (20 Hz), but the number of pulses varied (1, 3, 6, 11, 16, and 20 pulses) in 60% of total trials. Some mice started with variable pulse variants to counterbalance between subjects, and others began with the laser frequency variant (**Figure 4B**). In all experiments, the task lasted 30 min daily.

##### Frequency discrimination task

Mice were placed in an operant chamber (Med Associates Inc., VT, USA) equipped with three ports on one wall. The central port had a head entry detector (infrared sensor) and two lateral ports, and each contained a sipper that could deliver 2 µL dropper lick of 10% sucrose as a reward or a 10-psi air-puffs as a punishment. The task lasted for 30 min and consisted of delivering a laser stimulation after each head entry in the central port; the laser frequencies were either 10 or 20 Hz (1 s). Then, mice had to lick in the corresponding lateral port to receive 2 drops of sucrose. If mice licked in the incorrect port, they received 2 air-puffs. A clicker sound marked the start and the end of the trial (**Figure 5A**). A correct trial was when the mice licked in the port associated with the stimulation frequency. Mice learned the task when they reached 85% of correct trials in 3 consecutive sessions. In a fake laser session (**Figure 5D**), mice were connected to inactive optical fiber attached to the active fiber.

##### Frequency categorization task

Mice trained to discriminate between 10 vs. 20 Hz frequencies underwent a categorization task to indicate whether they received a lower or higher frequency. This task variant lasted 30 min, and it was similar to the frequency discrimination task, but one port was associated with lower frequencies (10, 12, and 14 Hz), whereas the opposite port corresponded to higher frequencies (16, 18, and 20 Hz).

##### Lever press self-stimulation task

Well-fed (sated) mice were placed in an operant chamber with two levers located in the front wall (Med Associates Inc., VT, USA). One lever was associated with the delivery of 1s of optogenetic stimulation (20 Hz), whereas the other was inactive (30 min daily sessions). After pressing the active lever, mice received a train of optogenetic stimulation (1 s “on” + 2 s of time out). After 3 sessions, the levers were switched to inactive, and thus the previously active lever was now inactive and vice versa. Mice were tested during 4 more sessions. In the extinction phase, the last 3 sessions, both levers were inactive.

##### Open field center self-stimulation

Sated animals were placed in a circular open field (diameter 45 cm) with a central circular flat glass (diameter 10 cm, thick 1.83 mm). In this task, mice had to cross to the central circular glass to receive optogenetic stimulation (1s, 20 Hz) + 2s of time out (**Figure 6D**) for 3 sessions (30 min each). In the last four sessions, optogenetic stimulation was not delivered as an extinction test.

##### Real-Time Conditioned Place Preference (rtCPP)

Well-fed mice were placed in a rectangular acrylic arena (20×20×42cm) that was divided in half with two different visual cues. One side had black/white stripes, and the other side had black/white circles. Mice were allowed to cross between sides freely. On day 1, mice explored the arena for 10 min (pre-test). On the following days (sessions 2 to 4), the 20 min consecutive sessions, they were conditioned on the side where they spent less time (or in the more preferred for LH^ArchT^ mice), by pairing optogenetic stimulation (20Hz, 1s “on”-2s “off” for ChR2 and eYFP, or continuous pulse 1s “on”+ 2s “off” for ArchT) each time they crossed and stayed in the less preferred side. On day 5, mice were placed in the arena again for 10 min without laser stimulation (test day). The preference index in the conditioned and unconditioned sides was computed by dividing the time spent on each side over the total exploration time (Gil-Lievana et al., 2020).

##### Open-loop task

Water-deprived LH^ArchT^ and LH^eYFP^ mice were placed in an operant chamber (Med Associates Inc., VT, USA), with a central sipper port to dispense 2 µL drops of sucrose 10% at each lick. They were stimulated in blocks of 1 min “laser on”-1 min ‘laser off’ during 30. All licking responses were recorded by a contact lickometer (Med Associates Inc., VT, USA).

##### Closed-loop task

Well-fed LH^ChR2^ and LH^eYFP^ control mice were placed in an operant chamber (Med Associates Inc., VT, USA), equipped with a central sipper port to dispense drops sucrose 10% each lick and a head entry detector (infrared sensor, Med Associates Inc., VT, USA). Each head entry triggered a train of optogenetic stimulation during 1 s (20 Hz) + 2 s time out in 30 min. Licks and head entries were recorded. Mice were placed in this task in 3 sessions, whereas the subsequent 3 were for extinction (without laser).

##### Histology

After experiments were finished, mice were treated with an overdose of pentobarbital sodium (0.1ml 10mg/Kg), and they were transcardially perfused with PBS followed by 4% paraformaldehyde (PFA). Brains were removed, stored for 1 day in PFA 4%, and later were exchanged with a 30% sucrose/PBS solution. Brains were sectioned at 40 µm coronal slices. Slices were placed in a mounting medium (Dako®) to visualize the fluorescence of the reporter (eYFP or tdTomato) and implantation sites. Images were taken with a Nikon eclipse e200 and with a progress gryphax microscope camera, using a 4x objective, and for visualization purposes, the image contrast was improved with Adobe Photoshop CS5.1 software.

## QUANTIFICATION AND STATISTICAL ANALYSIS

Data were analyzed in MATLAB R2021a (The MathWorks Inc., Natick, MA) and Graphpad Prism (La Jolla, CA, USA). Unless otherwise mentioned, data were expressed as a mean ± SEM, and statistical analysis was performed from a student’s t-test and two-way ANOVA test followed by a Holm-Sidak or Dunnett *post hoc* and repeated-measures ANOVA in case of comparison across sessions.

The firing rate was compared for electrophysiological recordings by a non-parametric Kruskal-Wallis test followed by a Tukey-Kramer *post hoc*. Neurons which firing rates during the laser “on” period were significantly different (*p*<0.05) at any laser frequency relative to the activity in the control trials were considered as modulated. The difference between firing rates during laser stimulation vs. control trials determined the modulation sign (i.e., increased or decreased).

## Supporting information

Movie S1. Optogenetic-cue alternation task, tone + laser. Example of transgenic mice in block 1

Movie S2. Only laser session. Transgenic mice in block 2, without tone

Movie S3. Fake laser session. Transgenic mice in block 3, without tone and with the flashing light of the laser

Movie S4. Only tone session. Transgenic mice without laser and with a tone as a cue.

Movie S5. Frequency discrimination task. Transgenic mice had to discriminate between two laser stimulation frequencies

Movie S6. Lever self-stimulation task. Example of NAcThy1 mice in a self-stimulation task

Movie S7. Open field center self-stimulation. Example of PFCThy1 mice crossing the center to self-stimulate

## Acknowledgments

We thank Professor Sidney A. Simon, Federico Bermudez-Rattoni, Pavel Rueda, and Román Rossi Pool for their thoughtful comments on this paper. To Mario Gil-Moreno for building homemade optrodes and Alam Coss, Raul Sanchez Granados, and Aketzali Garcia for technical help with experimental work. We also thank Ricardo Gaxiola, Victor Manuel Garcia Gomez, and Fabiola Hernandez Olvera for invaluable animal care.

## Funding

This project was supported in part by Productos Medix 3247 (R.G.), Cátedra Marcos Moshinsky (R.G.), fundación Miguel Aleman Valdes (R.G.).

## Author Contributions

J. LI and R.G. designed research. J. LI. performed research. J.LI. and M.L. performed rtCPP and frequency discrimination tasks. J. LI. and M.L. performed histological sections. J. LI analyzed data. B.F. and M.L. reviewed the paper. J. LI and R.G. wrote the paper

## Declaration of interests

Authors declare no competing interests

## Figures Supplements

**Figure 2--Supplement 1.**
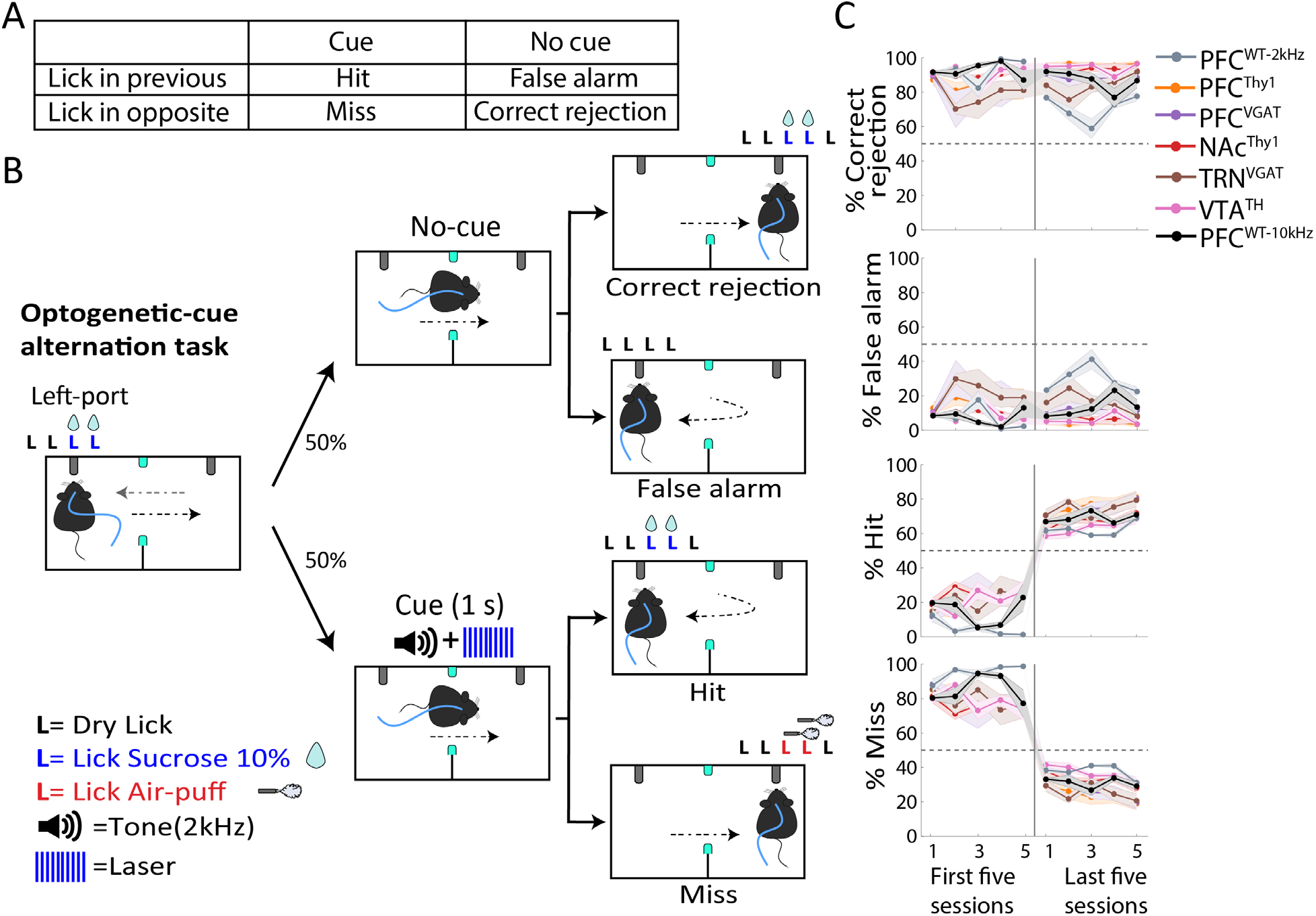
Mice learn to return to the previously rewarded port only when the optogenetic cue is present (Hit trials). (**A**) Table of different types of trials Correct Rejection, False Alarm, Hits, and Misses. In our task, mice had to continue alternating between sippers in no-cue trials (Correct Rejections), and very few False Alarms responses were observed. In contrast, in cue trials, lick responses were given in the previously rewarded sipper (Hits), and a few Misses were made. (**B**) Schematic of the optogenetic-cue sipper alternation task separated by trial type. (**C**) Performance in the first 5 and last 5 sessions during the task. After reaching the learning criteria (Horizontal dashed line), only hit trials increased while misses decreased. Correct rejection and False alarm maintained the same performance as in the initial first five sessions.

**Figure 2--Supplement 2.**
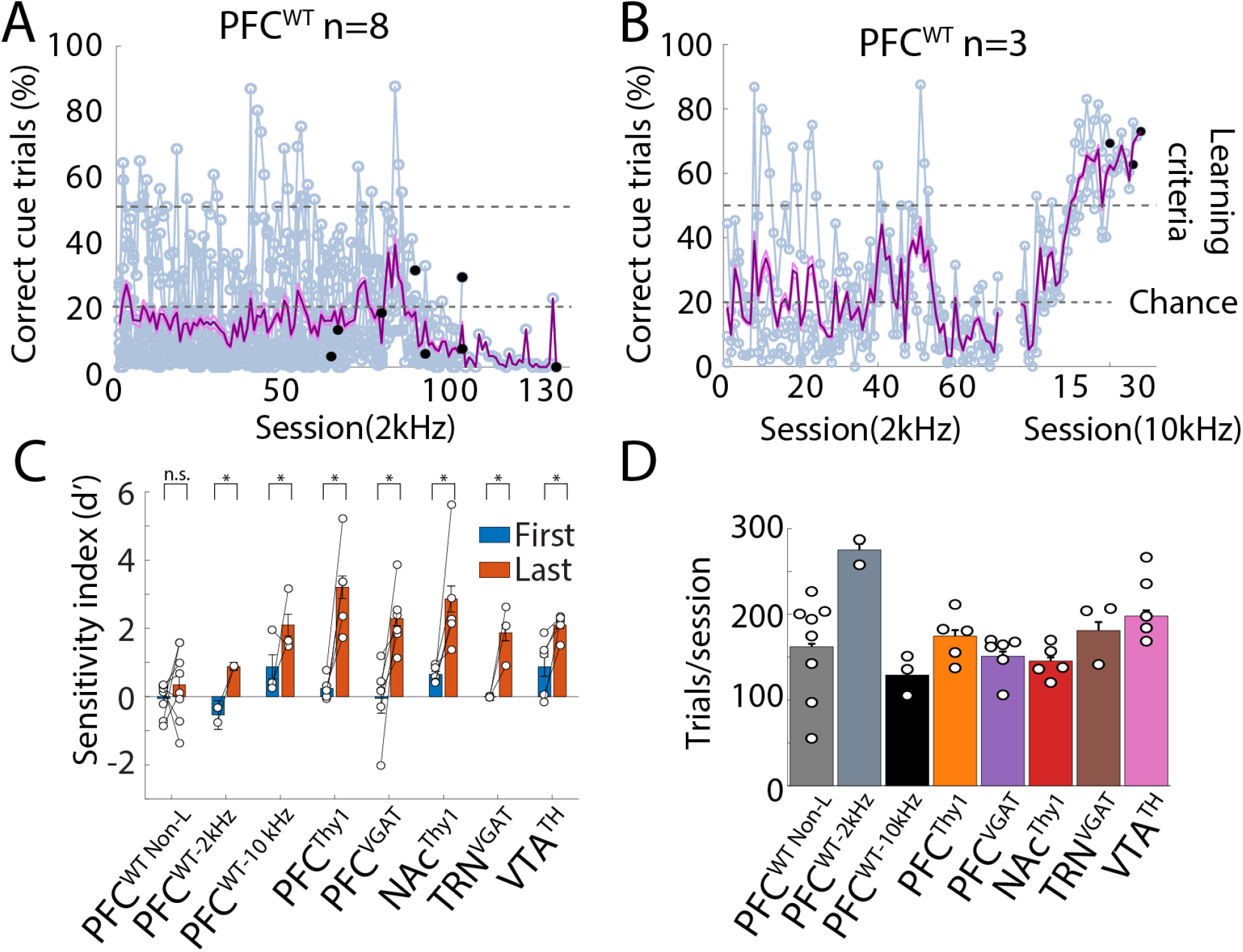
The 2 kHz tone was not as efficient as the optogenetic stimulation as a cue. (**A**) Correct trials of PFC^WT^ mice that did not reach the learning criteria trained at 2 kHz in the Optogenetic-cue sipper alternation task. Individual mice are shown in gray, and the mean ± SEM is shown in purple. Black dots showed the last session of each subject. (**B**) The tone frequency was changed to 10 kHz. The performance of the 3 non-learners subjects, from panel A. The same subjects were also trained with a 10 kHz tone. Note that after the tone was changed from 2 to 10 kHz, they rapidly learned the task (gray dash line learning criteria). (**C**) Sensitivity index (d prime, d’) computed for the first and last five sessions. Transgenic mice showed values above d’>1. In contrast, PFC^WT-2KHz^ exhibited values below d’<1, indicating that although they reached the learning criteria, they could not detect the cue as efficiently as transgenic mice or PFC^WT-10kHz^. PFC^WT Non-L^ refers to non-learners. (**D**) Total trials (Cue + No-Cue). Each dot represents an individual subject. \**p*<0.05 paired t-test.

**Figure 6—Supplement 1.**
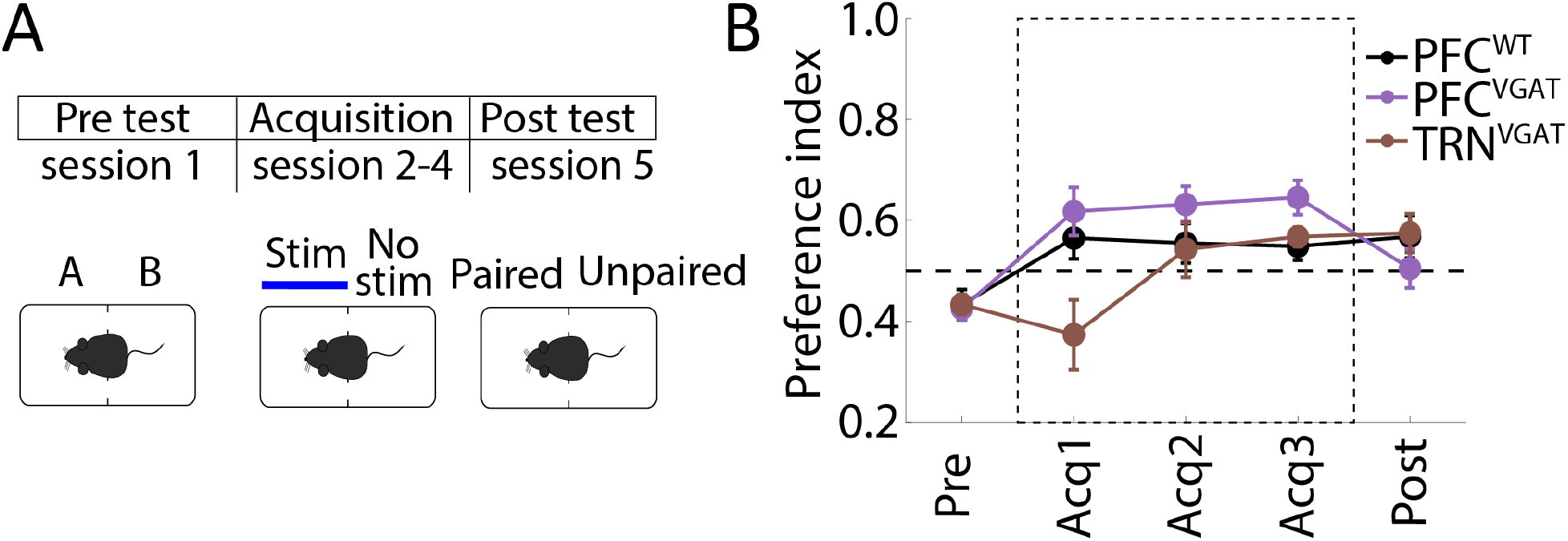
Activation of GABAergic neurons in PFC or TRN was not aversive neither rewarding. **(A)** Real-Time in Conditioned Place Preference task (rtCPP). Mice were placed in the box with two different contexts (A vs. B). Mice were stimulated in the less preferred side during 3 consecutive sessions, and finally, they were placed in a test session without stimulation. **(B)** Preference index in the side condition. Values above 0.5 mean that stimulation is preferred, while values below indicate that stimulation is avoided. PFC^VGAT^ and TRN^VGAT^ were not significantly different relative to PFC^WT^.

**Figure 7—Supplement 1.**
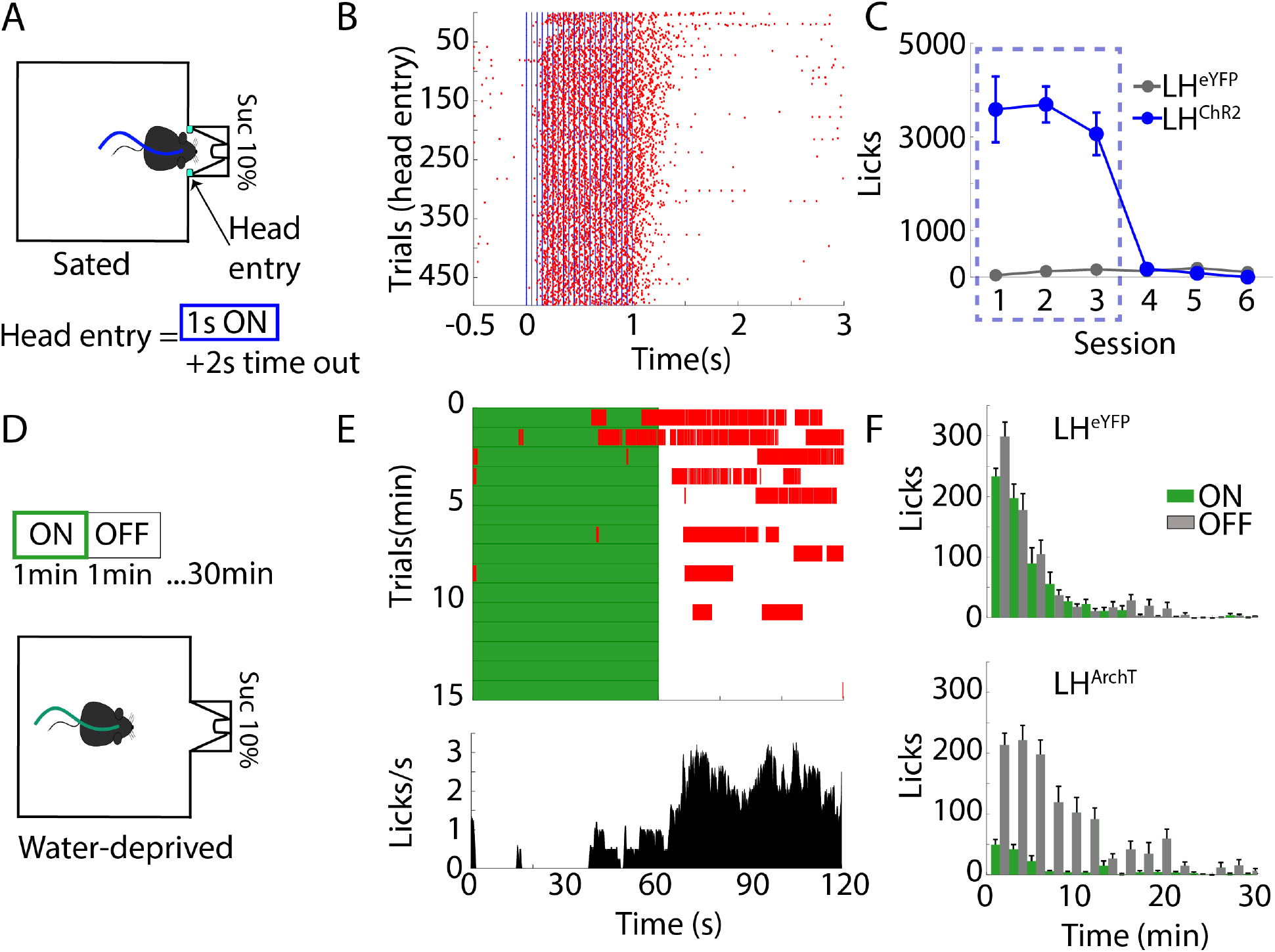
Activating or silencing of LH GABAergic neurons has opposing behavioral effects on feeding. (**A**) Schematic of closed-loop stimulation task. Sated LH^ChR2^ or LH^eYFP^ mice were placed in a behavioral box equipped with a sipper in a central port. The sipper was filled with sucrose 10%. In this task, the laser was triggered by a head entry in the central port (1 s, 20 Hz + 2 s time out, 473 nm). (**B**) Raster plot aligned to head entries for one LH^ChR2^ subject, red ticks = licks, blue ticks = laser. (**C**) Mean licks executed by LH^ChR2^ or LH^eYFP^ throughout sessions. The blue rectangle represents a laser session; the last three sessions were extinction sessions (no laser). (**D**) Schematics of the open-loop stimulation. Water-deprived LH^ArchT^ mice were located in a similar behavioral box than A, but with blocks of 1 min “on,” 1 min “off” (continuous pulse, at 532nm). (**E**) *Upper panel*, rater plot of one LH^ArchT^ mouse, aligned to laser onset (Time = 0), green rectangles indicate laser period, whereas red ticks indicate individual licks. Below is shown the PSTH average of lick responses across trials. (**F**) *Upper panel*, the histogram of each block of stimulation in the control LH^eYFP^ mice. Below, histogram of lick responses for LH^ArchT^ mice.

**Movie S1**.

Optogenetic-cue alternation task, tone + laser. Example of transgenic mice in block 1

**Movie S2**.

Only laser session. Transgenic mice in block 2, without tone

**Movie S3**.

Fake laser session. Transgenic mice in block 3, without tone and with the flashing light of the laser

**Movie S4**.

Only tone session. Transgenic mice without laser and with a tone as a cue.

**Movie S5**.

Frequency discrimination task. Transgenic mice had to discriminate between two laser stimulation frequencies

**Movie S6**.

Lever self-stimulation task. Example of NAc^Thy1^ mice in a self-stimulation task

**Movie S7**.

Open field center self-stimulation. Example of PFC^Thy1^ mice crossing the center to self-stimulate

## Notes

### Competing Interest Statement

The authors have declared no competing interest.

## References

Babl SS, Rummell BP, Sigurdsson T. 2019. The Spatial Extent of Optogenetic Silencing in Transgenic Mice Expressing Channelrhodopsin in Inhibitory Interneurons. Cell Reports 29:1381-1395.e4. doi:10.1016/j.celrep.2019.09.049

Berntson GG, Khalsa SS. 2021. Neural Circuits of Interoception. Trends in Neurosciences 44:17–28. doi:10.1016/j.tins.2020.09.011

Bevins RA, Besheer J. 2014. Interoception and Learning: Import to Understanding and Treating Diseases and Psychopathologies. ACS Chem Neurosci 5:624–631. doi:10.1021/cn5001028

Ceunen E, Vlaeyen JWS, Diest IV. 2016. On the Origin of Interoception. Frontiers in Psychology 7. doi:10.3389/fpsyg.2016.00743

Chow BY, Han X, Dobry AS, Qian X, Chuong AS, Li M, Henninger MA, Belfort GM, Lin Y, Monahan PE, Boyden ES. 2010. High-performance genetically targetable optical neural silencing by light-driven proton pumps. Nature 463:98–102. doi:10.1038/nature08652

Courtin J, Chaudun F, Rozeske RR, Karalis N, Gonzalez-Campo C, Wurtz H, Abdi A, Baufreton J, Bienvenu TCM, Herry C. 2014. Prefrontal parvalbumin interneurons shape neuronal activity to drive fear expression. Nature 505:92–96. doi:10.1038/nature12755

Craig AD. 2002. How do you feel? Interoception: the sense of the physiological condition of the body. Nat Rev Neurosci 3:655–666. doi:10.1038/nrn894

Danskin B, Denman D, Valley M, Ollerenshaw D, Williams D, Groblewski P, Reid C, Olsen S, Waters J. 2015. Optogenetics in Mice Performing a Visual Discrimination Task: Measurement and Suppression of Retinal Activation and the Resulting Behavioral Artifact. PLoS ONE 10:e0144760. doi:10.1371/journal.pone.0144760

Di Scala G, Mana MJ, Jacobs WJ, Phillips AG. 1987. Evidence of Pavlovian conditioned fear following electrical stimulation of the periaqueductal grey in the rat. Physiology & Behavior 40:55–63. doi:10.1016/0031-9384(87)90185-5

Doty RW. 1965. Conditioned reflexes elicited by electrical stimulation of the brain in macaques. Journal of Neurophysiology. doi:10.1152/jn.1965.28.4.623

Fetz EE. 1969. Operant Conditioning of Cortical Unit Activity. Science 163:955–958. doi:10.1126/science.163.3870.955

Fonseca E, de Lafuente V, Simon SA, Gutierrez R. 2018. Sucrose intensity coding and decision-making in rat gustatory cortices. eLife 7:e41152. doi:10.7554/eLife.41152

Garcia A, Coss A, Luis-Islas J, Puron-Sierra L, Luna M, Villavicencio M, Gutierrez R. 2021. Lateral Hypothalamic GABAergic Neurons Encode and Potentiate Sucrose’s Palatability. Front Neurosci 14. doi:10.3389/fnins.2020.608047

Gil-Lievana E, Balderas I, Moreno-Castilla P, Luis-Islas J, McDevitt RA, Tecuapetla F, Gutierrez R, Bonci A, Bermúdez-Rattoni F. 2020. Glutamatergic basolateral amygdala to anterior insular cortex circuitry maintains rewarding contextual memory. Commun Biol 3:139. doi:10.1038/s42003-020-0862-z

Guo W, Hight AE, Chen JX, Klapoetke NC, Hancock KE, Shinn-Cunningham BG, Boyden ES, Lee DJ, Polley DB. 2015. Hearing the light: neural and perceptual encoding of optogenetic stimulation in the central auditory pathway. Scientific Reports 5:10319. doi:10.1038/srep10319

Halassa MM, Chen Z, Wimmer RD, Brunetti PM, Zhao S, Zikopoulos B, Wang F, Brown EN, Wilson MA. 2014. State-Dependent Architecture of Thalamic Reticular Subnetworks. Cell 158:808–821. doi:10.1016/j.cell.2014.06.025

Heffner HE, Heffner RS. 2007. Hearing ranges of laboratory animals. J Am Assoc Lab Anim Sci 46:20–22.

Hölzl R, Erasmus L-P, Möltner A. 1996. Detection, discrimination and sensation of visceral stimuli. Biological Psychology 42:199–214. doi:10.1016/0301-0511(95)05155-4

Khalsa SS, Lapidus RC. 2016. Can Interoception Improve the Pragmatic Search for Biomarkers in Psychiatry? Front Psychiatry 7. doi:10.3389/fpsyt.2016.00121

Koralek AC, Jin X, Long Ii JD, Costa RM, Carmena JM. 2012. Corticostriatal plasticity is necessary for learning intentional neuroprosthetic skills. Nature 483:331–335. doi:10.1038/nature10845

Kumar S, Black SJ, Hultman R, Szabo ST, DeMaio KD, D. J, Katz BM, Feng G, Covington HE, Dzirasa K. 2013. Cortical Control of Affective Networks. Journal of Neuroscience 33:1116–1129. doi:10.1523/JNEUROSCI.0092-12.2013

Maffei A, Fontanini A. 2009. Network homeostasis: a matter of coordination. Curr Opin Neurobiol 19:168–173. doi:10.1016/j.conb.2009.05.012

Mazurek KA, Schieber MH. 2019. How is electrical stimulation of the brain experienced, and how can we tell? Selected considerations on sensorimotor function and speech. Cognitive Neuropsychology 36:103–116. doi:10.1080/02643294.2019.1609918

Mazurek KA, Schieber MH. 2017. Injecting Instructions into Premotor Cortex. Neuron 96:1282-1289.e4. doi:10.1016/j.neuron.2017.11.006

Nieh EH, Vander Weele CM, Matthews GA, Presbrey KN, Wichmann R, Leppla CA, Izadmehr EM, Tye KM. 2016. Inhibitory Input from the Lateral Hypothalamus to the Ventral Tegmental Area Disinhibits Dopamine Neurons and Promotes Behavioral Activation. Neuron 90:1286–1298. doi:10.1016/j.neuron.2016.04.035

Otchy TM, Wolff SBE, Rhee JY, Pehlevan C, Kawai R, Kempf A, Gobes SMH, Ölveczky BP. 2015. Acute off-target effects of neural circuit manipulations. Nature 528:358–363. doi:10.1038/nature16442

Owen SF, Liu MH, Kreitzer AC. 2019. Thermal constraints on in vivo optogenetic manipulations. Nature Neuroscience 22:1061–1065. doi:10.1038/s41593-019-0422-3

Penfield W, Rasmussen T. 1950. The Cerebral Cortex of Man: A Clinical Study of Localization of Function. New York: Macmillian.

Pinault D. 2004. The thalamic reticular nucleus: structure, function and concept. Brain Research Reviews 46:1–31. doi:10.1016/j.brainresrev.2004.04.008

Prado L, Luis-Islas J, Sandoval OI, Puron L, Gil MM, Luna A, Arias-García MA, Galarraga E, Simon SA, Gutierrez R. 2016. Activation of Glutamatergic Fibers in the Anterior NAc Shell Modulates Reward Activity in the aNAcSh, the Lateral Hypothalamus, and Medial Prefrontal Cortex and Transiently Stops Feeding. J Neurosci 36:12511–12529. doi:10.1523/JNEUROSCI.1605-16.2016

Prsa M, Galiñanes GL, Huber D. 2017. Rapid Integration of Artificial Sensory Feedback during Operant Conditioning of Motor Cortex Neurons. Neuron 93:929-939.e6. doi:10.1016/j.neuron.2017.01.023

Razran G. 1961. The observable and the inferable conscious in current Soviet psychophysiology: Interoceptive conditioning, semantic conditioning, and the orienting reflex. Psychological Review 68:81–147. doi:10.1037/h0039848

Romo R, Hernández A, Zainos A, Salinas E. 1998. Somatosensory discrimination based on cortical microstimulation. Nature 392:387–390. doi:10.1038/32891

Solinas M, Panlilio LV, Justinova Z, Yasar S, Goldberg SR. 2006. Using drug-discrimination techniques to study the abuse-related effects of psychoactive drugs in rats. Nature Protocols 1:1194–1206. doi:10.1038/nprot.2006.167

Sun Z, Schneider A, Alyahyay M, Karvat G, Diester I. 2021. Effects of Optogenetic Stimulation of Primary Somatosensory Cortex and Its Projections to Striatum on Vibrotactile Perception in Freely Moving Rats. eNeuro 8. doi:10.1523/ENEURO.0453-20.2021

Tabot GA, Dammann JF, Berg JA, Tenore FV, Boback JL, Vogelstein RJ, Bensmaia SJ. 2013. Restoring the sense of touch with a prosthetic hand through a brain interface. PNAS 110:18279–18284. doi:10.1073/pnas.1221113110

Verrier R, Calvert A, Lown B. 1975. Effect of posterior hypothalamic stimulation on ventricular fibrillation threshold. American Journal of Physiology-Legacy Content 228:923–927. doi:10.1152/ajplegacy.1975.228.3.923

Vong L, Ye C, Yang Z, Choi B, Chua S, Lowell BB. 2011. Leptin Action on GABAergic Neurons Prevents Obesity and Reduces Inhibitory Tone to POMC Neurons. Neuron 71:142–154. doi:10.1016/j.neuron.2011.05.028

Wimmer RD, Schmitt LI, Davidson TJ, Nakajima M, Deisseroth K, Halassa MM. 2015. Thalamic control of sensory selection in divided attention. Nature 526:705–709. doi:10.1038/nature15398

Wu Z, Zheng N, Zhang S, Zheng X, Gao L, Su L. 2016. Maze learning by a hybrid brain-computer system. Sci Rep 6:31746. doi:10.1038/srep31746

Yadav AP, Li S, Krucoff MO, Lebedev MA, Abd-El-Barr MM, Nicolelis MAL. 2021. Generating artificial sensations with spinal cord stimulation in primates and rodents. Brain Stimulation 14:825–836. doi:10.1016/j.brs.2021.04.024

Zhao S, Ting JT, Atallah HE, Qiu L, Tan J, Gloss B, Augustine GJ, Deisseroth K, Luo M, Graybiel AM, Feng G. 2011. Cell type–specific channelrhodopsin-2 transgenic mice for optogenetic dissection of neural circuitry function. Nature Methods 8:745–752. doi:10.1038/nmeth.1668

