## Supplementary figures for "Optoception: perception of optogenetic brain stimulation"

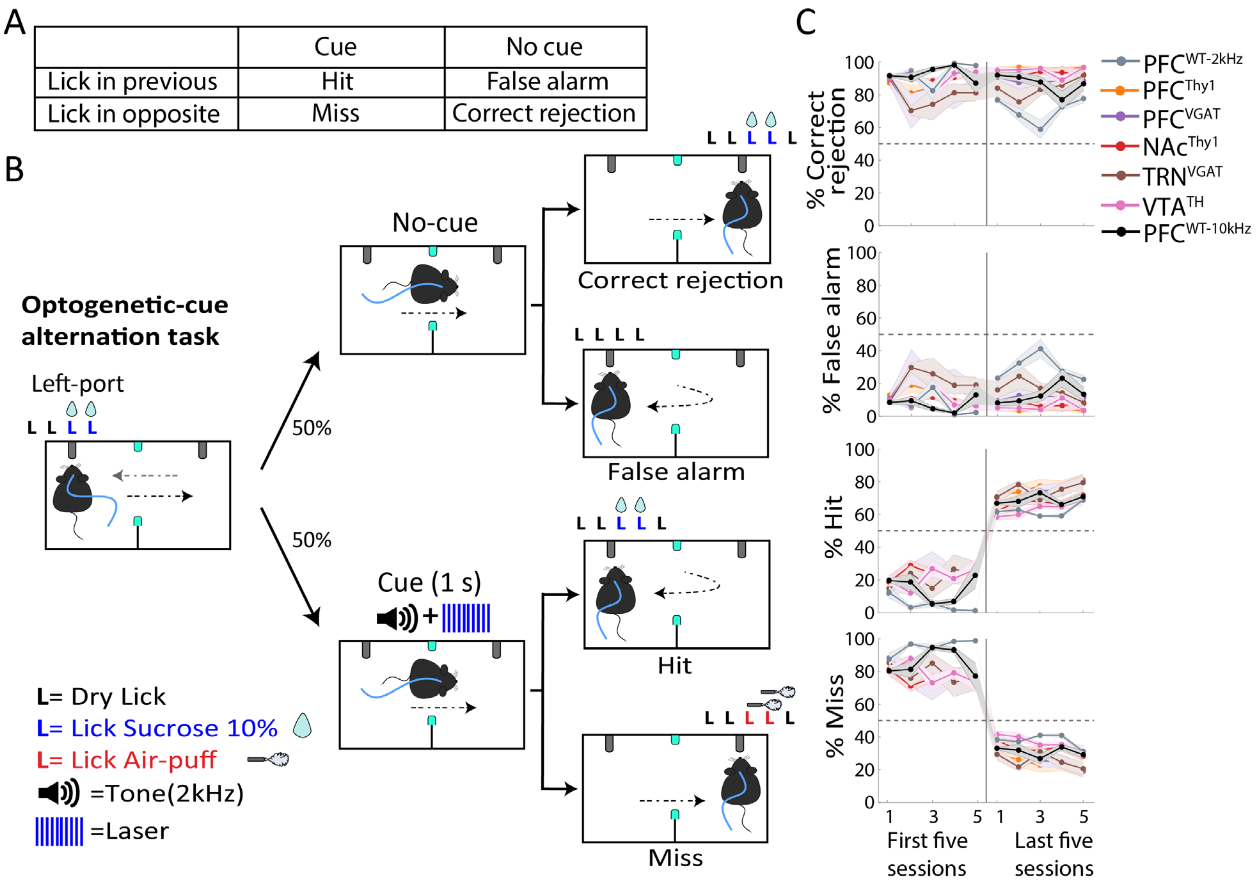


**Figure 2--Supplement 1. Mice learn to return to the previously rewarded port only when the optogenetic cue is present (Hit trials).**

(**A**) Table of different types of trials Correct Rejection, False Alarm, Hits, and Misses. In our task, mice had to continue alternating between sippers in no-cue trials (Correct Rejections), and very few False Alarms responses were observed. In contrast, in cue trials, lick responses were given in the previously rewarded sipper (Hits), and a few error Misses were observed. (**B**) Schematic of the optogenetic-cue sipper alternation task separated by trial type. (**C**) Performance in the first 5 and last 5 sessions during the task. After reaching the learning criteria (Horizontal dashed line), only hit trials increased while misses decreased. Correct rejection and False alarm maintained the same performance as in the initial first five sessions.

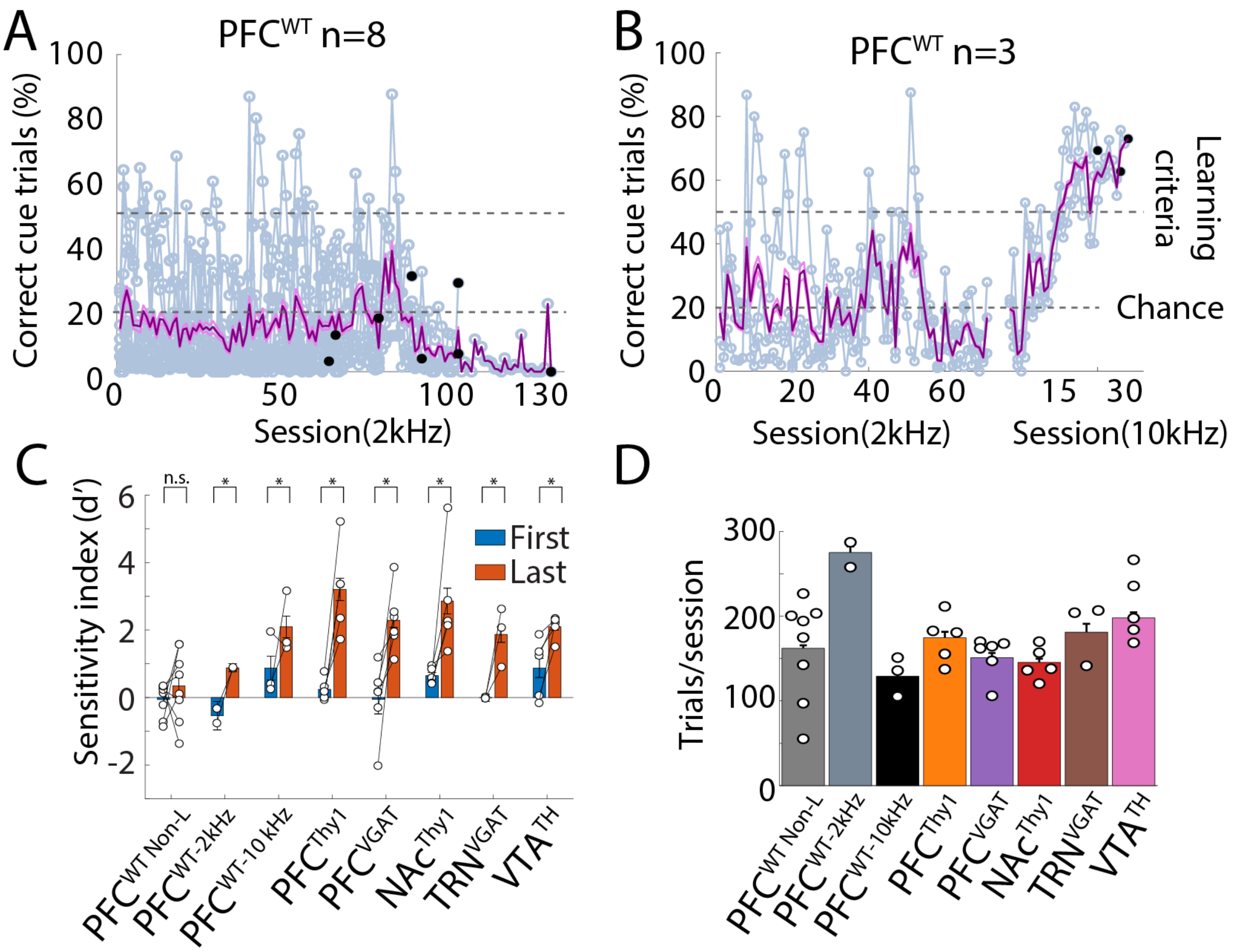


**Figure 2--Supplement 2. The 2 kHz tone was not as efficient as the optogenetic stimulation as a cue.**

(**A**) Correct trials of PFC$^{WT}$ mice that did not reach the learning criteria trained at 2 kHz in the Optogenetic-cue sipper alternation task. Individual mice are shown in gray, and the mean ± SEM is shown in purple. Black dots showed the last session of each subject. (**B**) The tone frequency was changed to 10 kHz. The performance of the 3 non-learners subjects, from panel A. The same subjects were also trained with a 10 kHz tone. Note that after the tone was changed from 2 to 10 kHz, they rapidly learned the task (gray dash line learning criteria). (**C**) Sensitivity index (d prime, d') computed for the first and last five sessions. Transgenic mice showed values above d'>1 while PFC$^{WT\text{-}2KHz}$ has shown values below d'<1, indicating that although they reached the learning criteria, they could not detect the cue as efficiently as transgenic mice or PFC$^{WT\text{-}10kHz}$. PFC$^{WT\ Non\text{-}L}$ refers to non-learners. (**D**) Total trials (Cue + No-Cue). Each dot represents an individual subject. *$p$<0.05 paired t-test.

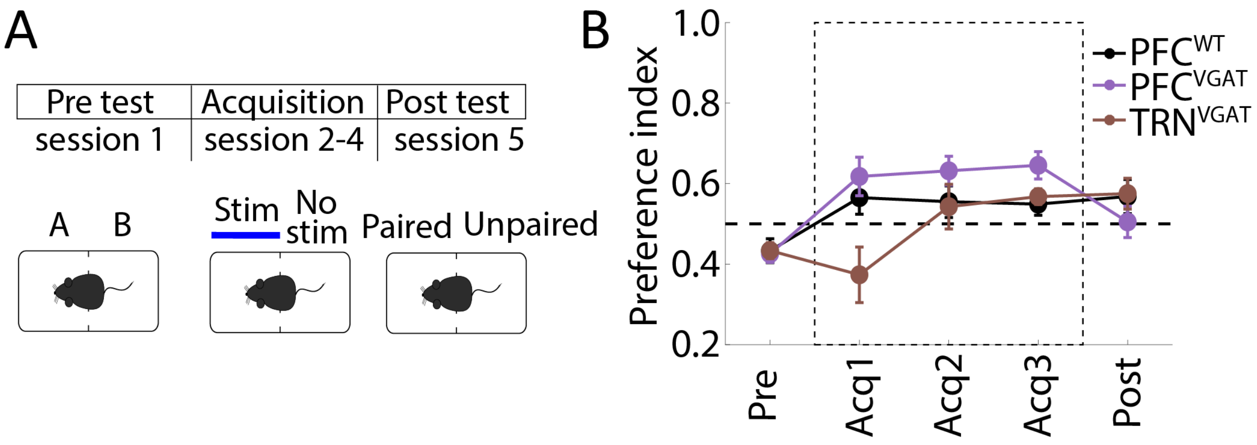


**Figure 6—Supplement 1. Activation of GABAergic neurons in PFC or TRN was not aversive neither rewarding. (A)** Real-Time in Conditioned Place Preference task (rtCPP). Mice were placed in the box with two different contexts (A vs. B). Mice were stimulated in the less preferred side during 3 consecutive sessions, and finally, they were placed in a test session without stimulation. **(B)** Preference index in the side condition. Values above 0.5 mean that stimulation is preferred, while values below indicate that stimulation is avoided. $PFC^{VGAT}$ and $TRN^{VGAT}$ were not significantly different relative to $PFC^{WT}$.

.

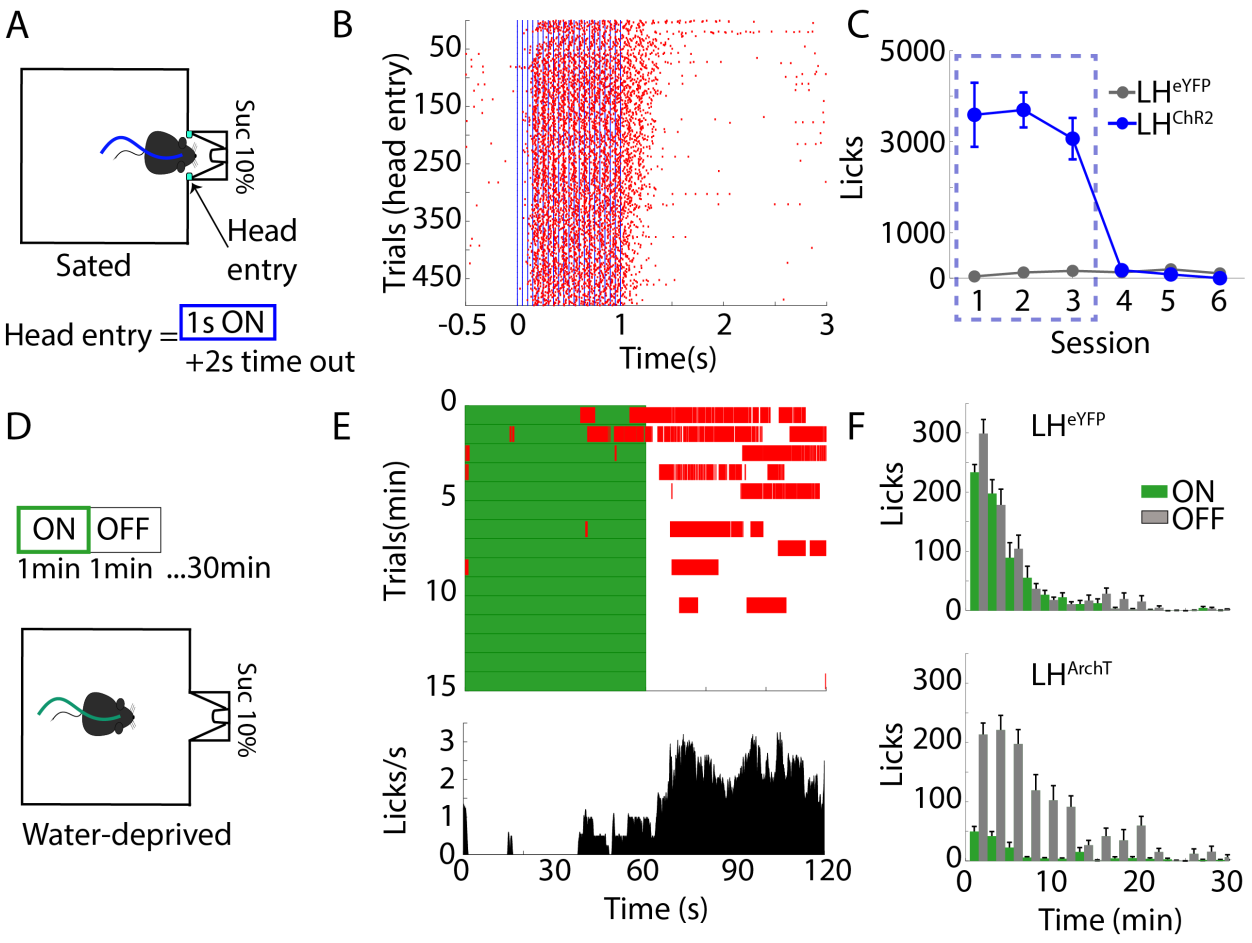


**Figure 7—Supplement 1. Activating or silencing of LH GABAergic neurons has opposing behavioral effects on feeding.** (**A**) Schematic of closed-loop stimulation task. Sated $LH^{ChR2}$ or $LH^{eYFP}$ mice were placed in a behavioral box equipped with a sipper in a central port. The sipper was filled with sucrose 10%. In this task, the laser was triggered by a head entry in the central port (1 s, 20 Hz + 2 s time out, 473 nm). (**B**) Raster plot aligned to head entries for one $LH^{ChR2}$ subject, red ticks = licks, blue ticks = laser. (**C**) Mean licks executed by $LH^{ChR2}$ or $LH^{eYFP}$ throughout sessions. The blue rectangle represents a laser session; the last three sessions were extinction sessions (no laser). (**D**) Schematics of the open-loop stimulation. Water-deprived $LH^{ArchT}$ mice were placed in a similar behavioral box than A, but with blocks of 1 min "on," 1 min "off" (continuous pulse, at 532nm). (**E**) *Upper panel*, rater plot of one $LH^{ArchT}$ mouse, aligned to laser onset (Time = 0), green rectangles indicate laser period, whereas red ticks indicate individual licks. Below is shown the PSTH average of lick responses across trials. (**F**) *Upper panel*, the histogram of each block of stimulation in the control $LH^{eYFP}$ mice. Below, histogram of lick responses for $LH^{ArchT}$ mice.

**Movie S1.**

Optogenetic-cue alternation task, tone + laser. Example of transgenic mice in block 1

**Movie S2.**

Only laser session. Transgenic mice in block 2, without tone

**Movie S3.**

Fake laser session. Transgenic mice in block 3, without tone and with the flashing light of the laser

**Movie S4.**

Only tone session. Transgenic mice without laser and with a tone as a cue.

**Movie S5.**

Frequency discrimination task. Transgenic mice had to discriminate between two laser stimulation frequencies

**Movie S6.**

Lever self-stimulation task. Example of $NAc^{Thy1}$ mice in a self-stimulation task

**Movie S7.**

Open field center self-stimulation. Example of $PFC^{Thy1}$ mice crossing the center to self-stimulate
